# Stage-dependent biotic interactions may not be important for stochastic competitive dynamics with little variation in stage structure

**DOI:** 10.64898/2026.03.13.711558

**Authors:** Young Jun Lee, Benjamin Wong Blonder, Courtenay A. Ray, Christina M. Hernández, Roberto Salguero-Gómez

## Abstract

1. Stage-dependent interactions, in which different life cycle stages (*e*.*g*., juveniles, adults) exert different per-capita competitive effects, are widespread across ecological communities. However, whether explicitly accounting for such ontogenetic variation improves forecasts of stochastic community dynamics remains unclear. We tested how the strength of stage dependence and species life-history strategy influence the predictive accuracy of community models that either include or ignore stage-specific interactions.
2. We constructed stochastic two-species competition models using stage-structured matrix population models spanning five virtual life histories along the fast-slow continuum. Density dependence was imposed separately on juvenile survival, adult survival, progression, retrogression, or fertility, and the strength of stage dependence varied from adult-driven to juvenile-driven competition. We then fitted deterministic projection models with and without stage-dependent interaction terms to simulated time series and quantified predictive performance over 100 time-step forecasts using mean absolute percentage error (MAPE).
3. Increasing stage dependence consistently reduced the predictive accuracy of models that ignored stage structure. However, absolute prediction errors remained small across all scenarios (MAPE < 0.7%), even under strong stage dependence. The influence of life-history strategy depended on which vital rate was density dependent: when juvenile survival was density dependent, faster life histories showed larger errors; when progression, retrogression, or fertility were density dependent, slower life histories exhibited greater errors; and when adult survival was density dependent, no consistent life-history effect emerged. Across simulations, temporal variation in population structure was low (coefficient of variation < 0.036), and prediction error was strongly associated with the magnitude of structural fluctuations rather than life-history pace per se.
4. *Synthesis*. Stage-dependent interactions can, in principle, alter stochastic competitive dynamics, but their practical importance for ecological forecasting depends on the extent to which population stage structure fluctuates through time. When environmental stochasticity dominates and stage structure remains near equilibrium, simpler models that ignore stage dependence provide robust approximations of community dynamics. Our results identify conditions under which demographic detail is necessary for forecasting and highlight the central role of structural variability in linking life-history strategy to community-level dynamics.

## Introduction

Biotic interactions are a major driver of the outcomes of community dynamics and coexistence. Indeed, these biotic interactions, the effects exerted by an organism on another, can lead to positive or negative consequences for the actor or recipient (Morales-Castilla et al., 2015). These interspecific interactions manifest as density-dependent effects, where the net biotic interaction experienced by a recipient varies with the density of the actors (Murray, 1982). Density-dependent effects may influence species’ population growth rates (Berryman et al., 2002; Murray, 1982) and underlying vital rates (*i*.*e*., survival, growth, reproduction; Neubert & Caswell, 2000). In turn, density-dependent effects can drive the dynamics of communities, together with environmental and demographic stochasticity (Mutshinda et al., 2009). Particularly, density-dependent effects underlie the regulation of communities by competition (Hanski, 1997; Peters, 2003) or trophic interactions (Heath et al., 2014; Sibly et al., 2002; Thakur & Geisen, 2019).

Anthropogenic drivers such as climate change, biological invasions, habitat fragmentation and harvesting can alter both the strength and the stage-specific expression of biotic interactions. These drivers can potentially disrupt the density-dependent mechanisms that regulate communities and mediate coexistence (Blois et al., 2013; Waller et al., 2020). For example, warming can shift phenology and ontogenetic timing (Pausas 2025), invasive species can differentially affect juvenile and adult stages (McCard et al., 2024), and selective harvesting or disturbance can skew population stage structures (Matte et al., 2023). Such changes may modify not only overall interaction strength but also how interaction effects are distributed across life stages, thereby altering population growth rates, competitive outcomes and community stability. Because density-dependent feedbacks operate through vital rates, and because these vital rates often vary across ontogeny (Caswell 2001), anthropogenic perturbations may generate community responses that depend critically on demographic structure. In this context, community models that explicitly represent biotic interactions and potentially their stage dependence are increasingly viewed as essential tools for conservation and management, enabling forecasts of community trajectories under alternative environmental and management scenarios (Boult, 2023; Petchey et al., 2015). However, the level of demographic detail required for reliable forecasts remains uncertain, particularly under stochastic environmental change.

Biotic interactions are shaped by the ontogenetic stage of the organisms involved in them. Ontogenetic niche shift refers to a shift in individual traits and thus abiotic and biotic interactions along an ontogenetic dimension (*e*.*g*., age, size, *etc*.; Parish & Bazzaz, 1985). In this context, a key aspect of ontogenetic niche shift is ontogenetic variation in biotic interactions (Miller & Rudolf, 2011), whereby the magnitude and direction of density-dependent effects can vary within a population depending on the life stage of interacting individuals. For example, adults of the desert sagebrush (*Ambrosia dumosa*) promote conspecific juvenile survival but hinder conspecific adult growth via competition (Miriti, 2006). Stage-dependent interactions are widespread. Examples can be found in the trophic ecology of aquatic animals (Persson & Crowder, 1998; Rudolf, 2020) and competition among insect herbivores (Karban, 1989) and between herbaceous plants (Grace, 1985; Leger & Espeland, 2010; Schiffers & Tielbörger, 2006). However, because different life stages can exert opposing or asymmetric per-capita effects, the net population-level consequence of stage-dependent interactions is not immediately apparent and may depend critically on the relative abundance of stages within each population.

When biotic interactions are stage-dependent, population structures also can modulate density-dependent effects. These effects can then impact community dynamics (Miller & Rudolf, 2011; Fig. 1A & 1B). Prior work has focused primarily on deterministic models, suggesting that the influence of population structure on density-dependent interactions can have community-level consequences (Miller & Rudolf, 2011). Examples include increased community stability (de Roos, 2021; Romero et al., 2025), emergence of alternative stable states (Nakazawa, 2011a, 2011b), and alternative conditions for competitive coexistence with bottom-up (Loreau & Ebenhoh, 1994; Moll & Brown, 2008; Schellekens et al., 2010) or top-down regulation (Wollrab et al., 2013). In sum, stage-dependent interactions introduce population structure as an additional dimension controlling density-dependent effects alongside population abundance because the population-average per-capita interaction strength emerges from the combination of stage-specific effects and the relative abundance of the stages (Miller & Rudolf, 2011; Nakazawa, 2015).

**Figure 1.**
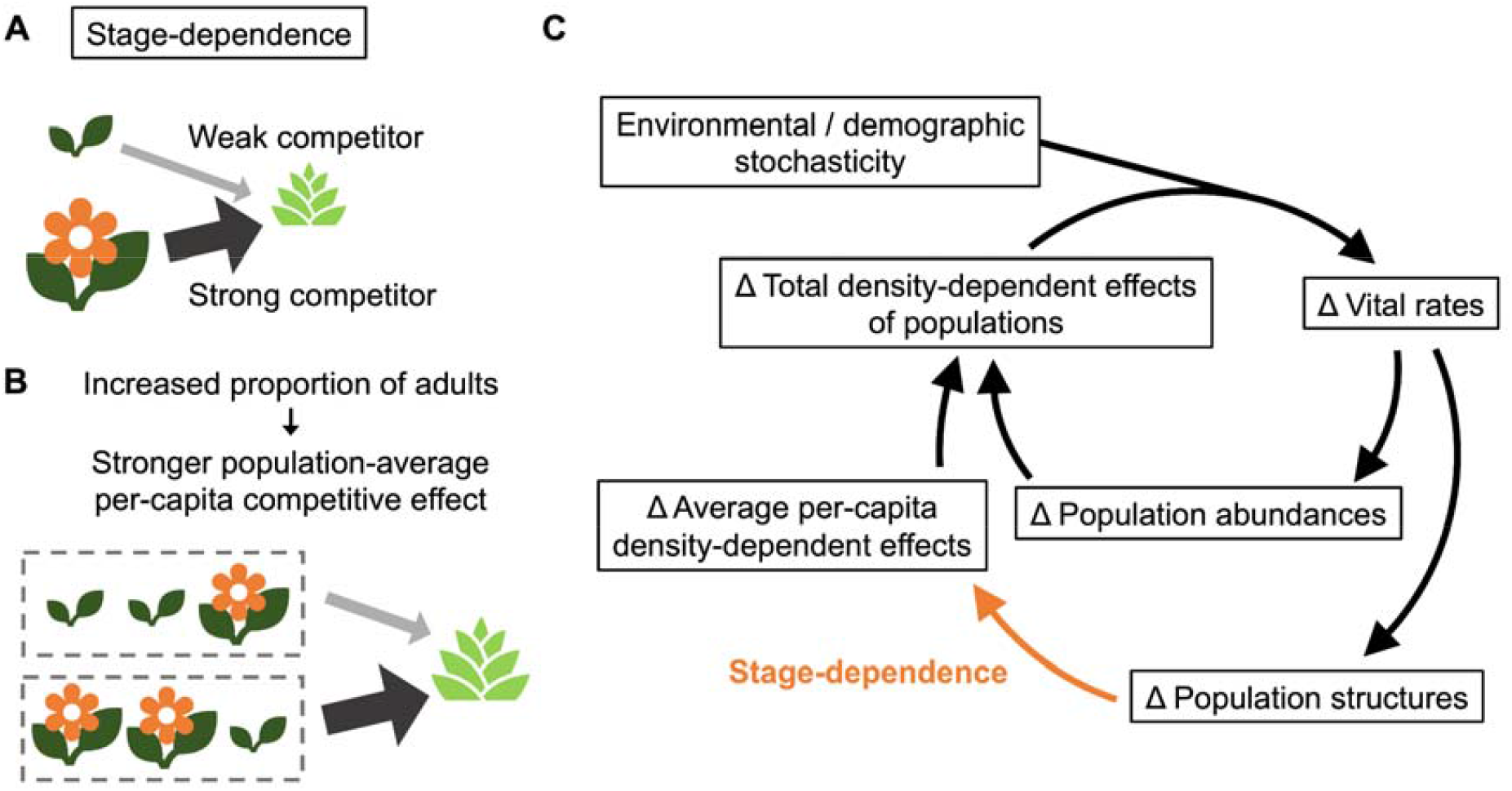
Stage-dependence in biotic interactions causes population-average per-capita interactions to depend on the structure of the interacting species’ populations, which in turn may impact the dynamics of the whole community. (**A**) A hypothetical example of stage-dependent interactions. Here, the per-capita competitive effect exerted by adult plants of a species on another species is greater than that of juvenile plants of the first species. (**B**) In turn, stage-dependent competition generates an association between a population’s structure and its average per-capita competitive effect. In this example, an increase in the proportional abundance of the more competitive adults increases the average per-capita competitive effect of the population of said species. (**C**) The potential role of stage-dependent interactions in modulating the stochastic dynamics of the whole community. Environmental and demographic stochasticity alter population-level vital rates of survival, growth, and reproduction, which generate fluctuations in population abundances and structures of the species in a given community. Then, population abundances and structures modulate the magnitude and direction of population-level density-dependent effects. Notably, stage-dependent interactions allow population structure to influence population-level density-dependent effects.

The effect of stage-dependent interactions on the importance of population structure in modelling stochastic community dynamics remains unknown. Natural communities are constantly exposed to environmental and demographic stochasticity (Crone, 2016; Sæther, 1997) and to aperiodic disturbances (Bazzaz, 1983). Stochasticity and disturbances cause variability in population abundance and structure, which in turn can alter vital rates via density dependence (Sinclair & Pech, 1996; Fig. 1C). Correspondingly, the importance of population structure in driving fluctuations in population abundances may increase with (1) stronger stage dependence (*i*.*e*., greater between-stage difference between per-capita interaction strengths) and (2) greater variation in population structure. Of these factors, variation in population structure depends on the focal species’ life history strategy (Capdevila et al., 2022; Sæther et al., 2013) (*i*.*e*., collection of traits describing an organism’s life cycle *sensu* Stearns (1992)). Indeed, some evidence exists that where population structure is a significant driver of community dynamics due to these factors, a community forecast model without stage-dependent interactions may produce inaccurate predictions (Miller & Rudolf, 2011).

It remains unclear how important it is to account for stage dependent interactions when assessing community level outcomes. On the one hand, parameterising a complex community model with stage-specific interactions would likely demand data with greater resolution and volume (Getz et al., 2018). These data may not be available due to limited time and resources in a conservation context (Geary et al., 2020). On the other hand, given the large stochasticity in nature and the clear biological importance of stage variation, simplistic models may miss important components of the complex interactions they aim to describe. Given this trade-off, identifying the conditions under which such a complex model is required *vs*. conditions where a simpler model is adequate is critical for both theoretical and empirical applications.

Here, we explore the impact of stage-dependent interactions on the importance of population structure in modelling stochastic dynamics (Fig. 2). Specifically, we ask how stage-dependence in biotic interactions affect the predictive accuracy of community models with or without stage dependence. To address this question, we simulate stochastic two-species competition models with varying stage-dependent interactions. To do so, we use virtual species with linearly interpolated vital rates between extremes of the fast-slow continuum (Gaillard et al., 1989; Salguero-Gómez et al., 2016). A faster life history is characterised by shorter generation time (*i*.*e*., average age of reproduction), lower juvenile survival, faster growth, and greater reproduction (Gaillard et al., 1989; Salguero-Gómez et al., 2016). Then, on the simulated time series, we parameterise deterministic stage-structured community models with or without stage-dependent interactions. We explore two key hypotheses whose testing would provide general insights into these questions. We expect that (**H1**) increased stage dependence in biotic interactions will reduce the predictive accuracy of the community model without stage-dependent interactions. This is because when per-capita interaction strengths differ among life stages, the population-average density-dependent effect varies with stage structure; models that ignore stage dependence assume a constant interaction strength and thus miss variation driven by demographic shifts. We also expect that (**H2**) the effect of altered stage dependence will be stronger when both the focal species and its competitor have faster life histories. This is because faster life histories are generally associated with greater demographic variability (Sæther et al., 2013) and greater variation in transient population growth rate (Capdevila et al., 2022).

**Figure 2.**
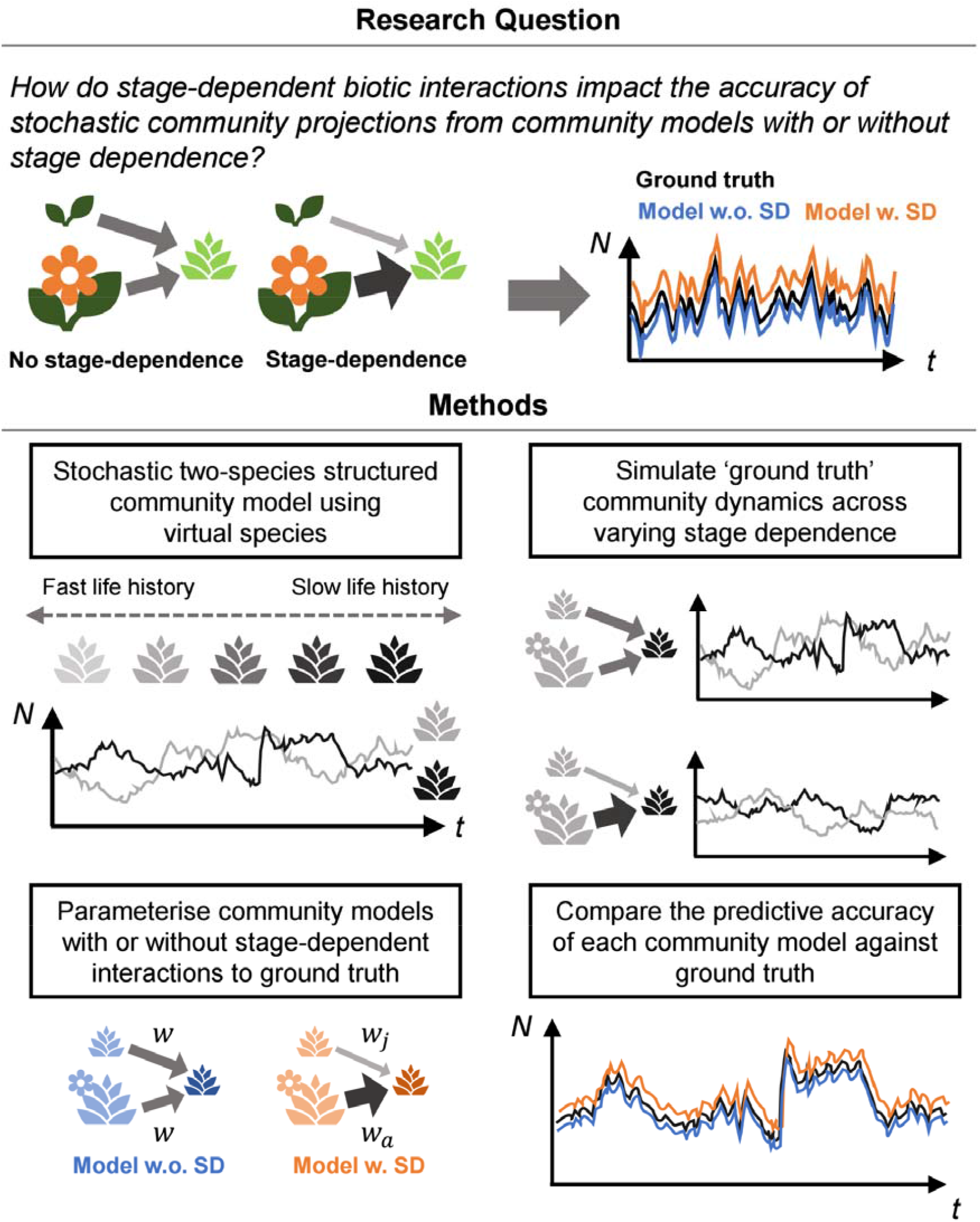
An overview of the questions and methods of this study. (**Top**) Here, we ask how stage-dependence in biotic interactions affects the predictive accuracy of community models with or without stage dependence. (**Bottom**) We construct a stochastic competition model for two virtual species with density-dependent effects and environmental stochasticity on vital rates. The virtual species span the fast-slow continuum of life history strategies. We vary the strength of stage dependence in density-dependent effects and simulate each community through time. Then, to the time series data, we parameterise community projection models with or without stage-dependent biotic interactions and compare their predictive accuracy.

## Materials and methods

Here, we employ a theoretical approach in which we quantify the potential impact of stage-dependent interactions on the accuracy of community models with or without stage dependence to test the aforementioned hypotheses.

### Model construction

To simulate stochastic community dynamics, we constructed a structured competition model for two virtual species. We modelled the population of each of the two species using a matrix population model (MPM), which describes discrete-time transitions and per-capita contributions across discrete life stages (Caswell, 2001). Our two-stage model follows Hernández et al. (2026), such that we projected the two-stage MPM for each species ***i*** from year ***t*** to ***t* + 1**, thus explicitly accounting for the effects of juvenile survival (***σ***_***j***,***i***_**(*t*)**), adult survival (***σ***_***o***,***i***_**(*t*)**), progression from juvenile to adult (***γ***_***i***_**(*t*)**), retrogression from adult to juvenile (***ρ***_***i***_**(*t*)**), and fertility (***ϕ***_***i***_**(*t*)**) as follows:

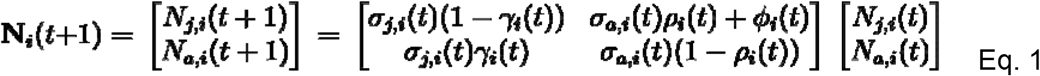

Here,**N**_***i***_**(*t*+1**) is the population vector of species ***i*** in year ***t* + 1**, while ***N***_***j***,***i***_**(*t*)** and ***N***_***a***,***i***_**(*t*)** are stage-specific abundances of juveniles and adults in year ***t***, respectively. This model assumes that progression and retrogression occur shortly before the census, while reproduction occurs shortly after the census (*i*.*e*., pre-breeding model design).

To generate a community time series with competition and stochasticity, we modelled vital rates as random variables controlled by density-dependent effects from both conspecific and heterospecific densities. Specifically, we modelled the density-dependent vital rate of each species ***i*** in year ***t***(***θ***_***i***_**(*t*)**) as a transformed sum of an environmental stochasticity term ***ϵ***_***θ***,***i***_**(*t*)**, a baseline vital rate term, ***c***_***θ***,***i***_ and a density-dependence term ***δ***_***θ i***_**(*t*)** as follows:

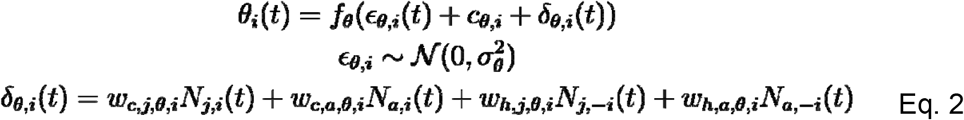

Here, ***f***_***θ***_**(*x*)** is the vital rate-specific transformation function (Eqs. 3 and 4), 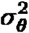 is the variance of the stochasticity term, and ***w***_***c***,***s***,***θ***,***i***_ and ***w***_***h***,***s***,***θ***,***i***_ are weights controlling the per-capita density-dependent effect of individuals in stage ***s*** (juvenile (***j***) or adult (***a***)) in the same species (subscript *c* for conspecifics) or the other species (subscript *h* for heterospecifics) in this two-species virtual community, respectively. When referring to elements of the population density vector ***N***, we use the species subscript −***i*** to refer to the competitor of species ***i***. To examine the effect of stage-dependent competition on one vital rate at a time, we modelled all vital rates except the focal vital rate as a constant, which we refer to as 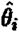.

To restrict each vital rate to biologically and mathematically valid ranges, we used bounded identity functions as transformation functions. Specifically, we used

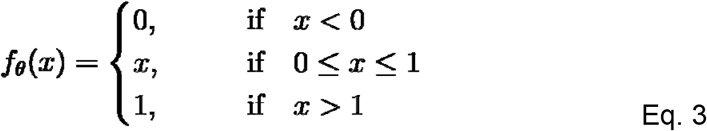

to restrict ***σ***_***j***_, ***σ***_***a***_, ***γ***, and ***ρ*** to [0,1], as these vital rates correspond to probabilities, and used

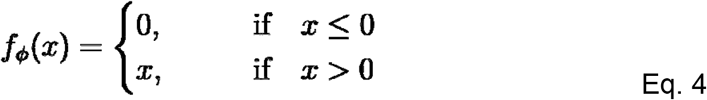

to restrict ***ϕ*** to **[0,+∞]**, as reproduction is constrained between it not happening (0) and potentially taking place at high positive values.

We modelled stochasticity and density dependence as having linear and additive effects in Eq. 2 to aid interpretation of results in relation to life history strategy. Previous density-dependent MPMs have typically scaled vital rates with a negative exponential function of population size (Jensen, 1995; Neubert & Caswell, 2000; Smouse & Weiss, 1975) or a nonlinear function analogous to a logistic growth model (Miller et al., 2002). Although these approaches prevent vital rates from taking unrealistic values, the methods cause the rate of change of vital rates with density to depend on the chosen baseline value (*e*.*g*., the value of the vital rate at approximately zero density of competitors). As such, interpreting community dynamics as the joint effect of fluctuating densities and life-history-dependent metrics (*e*.*g*., sensitivity of asymptotic growth rate to changes in vital rates) may be less intuitive. To ensure that per-capita density-dependent effects depend only on the strength of competition (*i*.*e*., ***w***_***c***,***s***,***θ***,***i &***_ ***w***_***h***,***s***,***θ***,***i***_), we modelled density dependence as an additive term and used linear transformation functions. Following the same logic, we also modelled stochasticity as an additive term, which is analogous to some previous stochastic MPMs (Miller et al., 2002; Sykes, 1969). Below, we further describe adjustments to density-dependent weights and stochastic variance to bound vital rates within the linear portion of the transformation function (Eq. 3-4).

### Model parametrisation

To generate simulated community time series data with varying levels of stage dependence, we defined baseline weights of density-dependent interactions (Eq. 5) that satisfy several criteria. First, we defined the “baseline weight” as the strength of competition in the absence of stage dependence. Therefore, we set the baseline weights of juveniles and adults of the same species to be equal. Second, to ensure that density-dependent effects on population growth rate are negative, we set the sign of the weights to the inverse of the sensitivity of population growth rate to the density-dependent vital rate (Supplementary Information S1). In practice, this means that competition decreases all vital rates except for retrogression, which increases in high-competition environments (see Hernández et al. 2026 and citations therein). Third, to control the relative strength of intraspecific and interspecific interactions, we set conspecific weights to be 4.5 times greater than the corresponding heterospecific weights. This ratio reflects a recent meta-analysis suggesting that intraspecific competition is, on average, 4-5 times stronger than interspecific competition in plant communities (Adler et al., 2018). Fourth, for symmetric competition, we set baseline weights to be constant across simulations with a particular density-dependent vital rate. In other words, the life histories of the focal species and the competitor species did not influence the baseline weights and instead the baseline weights are dependent on minimum values of vital rates across all virtual species. Finally, to restrict vital rates within the linear portion of the transformation function (Eqs. 3-4), we scaled the weights with the minimum baseline value of the density-dependent vital rate across the virtual life histories.

In sum, we defined the baseline effects of the density of conspecifics or heterospecifics of life stage ***s*** on the focal vital rate ***θ*** as follows:

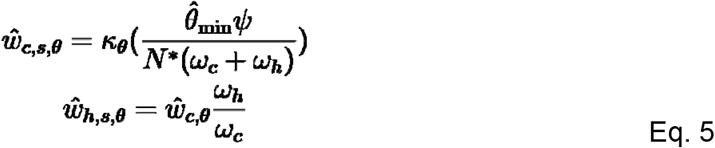

The descriptions and values of each coefficient in Eq. 5 are summarised in Table 1. Because baseline fertilities depend on the baseline interaction weights (see Virtual life histories, below), we assigned an arbitrary value of 0.1 to 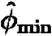. See Table S1 for details regarding the baseline interaction weights for each density-dependent vital rate.

**Table 1.** Summary of the parameters used to calculate the baseline biotic interaction effects used across simulations. The description and value of each parameter is provided.

| Parameter | Description | Value |
| --- | --- | --- |
| $\omega_c$ | Coefficient of the strength of conspecific interactions relative to heterospecific interactions | 1 |
| $\omega_h$ | Coefficient of the strength of heterospecific interactions relative to conspecific interactions | $\frac{1}{4.5}$ |
| $\hat{\theta}_{min}$ | Minimum of the baseline values of focal vital rate □ across virtual life histories | 0.1 for fertility;<br>See Table 2 for other vital-rate-specific values |
| $\psi$ | Constant scaling factor for interaction weights according to the minimum vital rate | 0.3 |
| $N^*$ | Target population size at equilibrium | 500 |
| $\kappa_\theta$ | Directional effect of increased density on the focal vital rate | −1 for juvenile survival, adult survival, progression, and fertility;<br>1 for retrogression |

To manipulate the degree of stage-dependence without altering the overall strength of competition we introduced a tunable stage-dependence factor. This factor, ***r***_**Δ**_, redistributes per-capita interaction strength between life stages while holding the population-average interaction strength constant at equilibrium. Specifically, ***r***_**Δ**_ controls the degree of stage dependence across all interactions in a simulation with density-dependent vital rate:

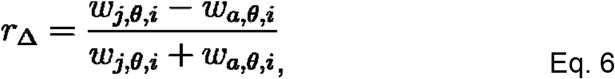

where the possible values of ***r***_**Δ**_ range from −1 (*i*.*e*., only adults exert density-dependent effects), through 0 (*i*.*e*., per-capita density-dependent effects of juveniles and adults are equal), to 1 (*i*.*e*., only juveniles exert density-dependent effects). Note that ***r***_**Δ**_ is equivalent for conspecifics and heterospecifics.

In manipulating stage-dependent interactions, we constrained the population-average per-capita interaction strengths to remain constant at stationary equilibrium population structures. That is, for a given equilibrium proportion of juveniles and adults in the population of species ***i*** (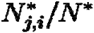 and 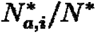, respectively) the stage-dependent weights must relate to the baseline weight as follows:

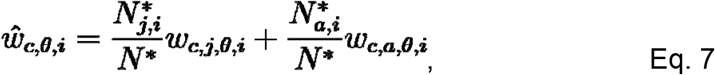

with the same constraint holding for the heterospecific weights.

To numerically determine stationary equilibrium stage structures, we simulated the community with baseline weights for each species pair and density-dependent vital rate. Then, we calculated the mean stage structures of each population from ***t*** = 1,000 to ***t*** = 2,000 time steps, and calculated the average of those stage structures across 10 replicate simulations. To obtain the final stage-specific density-dependence weights at a given value of ***r***_**Δ**_, we solved Eqs. 6 and 7 simultaneously.

To bound vital rates within the linear portion of the transformation function, we scaled the strength of stochasticity with the minimum value of each vital rate. Specifically, we set the standard deviation of the stochasticity term (***σ***_***θ***_) to 10% of the minimum vital rate as given in Table 2. As a result, the effects of density and stochasticity terms on vital rates were largely linear across all simulations.

**Table 2.** Summary of the baseline vital rates of the five virtual life history strategies used here to address hypothesis H2 on the interaction between stage-dependent interactions and life history. F(ast)2 represents the fastest life history strategy, followed by F1, M(edium), S(low)1, and S2 in descending pace of life.

| **Baseline vital rate** | **Life history strategy** |  |  |  |  |
| --- | --- | --- | --- | --- | --- |
|  | **F2** | **F1** | **M** | **S1** | **S2** |
| Juvenile survival | 0.400 | 0.525 | 0.650 | 0.775 | 0.900 |
| Adult survival | 0.600 | 0.688 | 0.775 | 0.862 | 0.950 |
| Progression | 0.900 | 0.750 | 0.600 | 0.450 | 0.300 |
| Retrogression | 0.050 | 0.062 | 0.075 | 0.088 | 0.100 |
| Minimum fertility | 1.132 | 0.754 | 0.490 | 0.283 | 0.109 |
| Maximum fertility | 1.629 | 1.125 | 0.809 | 0.561 | 0.344 |

### Virtual species

To test hypothesis H2 on the joint effect of stage dependence and life history, we defined five virtual life history strategies spanning the fast-slow continuum (Salguero-Gómez et al., 2016; Stearns, 1992). In the present model, we represented faster life histories with lower juvenile (***σ***_***j***_) and adult survival (***σ***_***a***_) and retrogression (***ρ***) and higher progression (***γ***) and fertility (***ϕ***) (Table 2). To restrict baseline vital rates to realistic values, we bounded the four vital rates, excluding fertility, within the virtual species range selected by Hernández et al. (2026) based on empirical models the COMPADRE plant MPM database (Salguero-Gómez et al., 2015; reported by Hernandez et al., 2026).

To test H2 in the absence of confounding effects, we calculated fertility to control equilibrium population sizes (Table 2). Specifically, we calculated fertility such that when both populations are at the target equilibrium size ***N*\*** = 500, the asymptotic growth rate (**λ**) is equal to 1. As such, we derived an expression for **λ** in terms of vital rates and solved **λ** = 1 for fertility when the density-dependent vital rate ***θ*** is equal to:

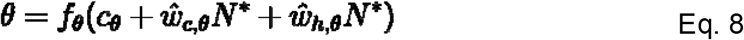

We provide a detailed account of this procedure in Supplementary information S2. Indeed, long-term mean population sizes were controlled between 490 and 510 across all simulations (Fig. S1).

To verify that these life histories have the desired demographic properties, we constructed MPMs from the baseline vital rates. From the MPMs, we calculated generation time (*i*.*e*., average age of reproductive individuals in the population) and life expectancy using age-from-stage decomposition methods for matrix population models described in Caswell (2001, pp. 110-132). Indeed, the modelled fastest life history was associated with the shortest generation time (3.2 ~ 3.4 years) and life expectancy (2.0 years). In contrast, the slowest life history had the longest generation time (8.0 ~ 9.5 years) and life expectancy (14.8 years). This pattern was preserved with the mean realised vital rates after density-dependent and stochastic effects (Table S2).

### Quantifying response in stochastic community dynamics

Because we are interested in how the effects of stage-dependence on stochastic community dynamics differ based on the life history strategy of the focal species, the life history of the competitor, and the vital rate that is subject to density-dependent competition, we designed a series of simulations with a fully factorial set of possible combinations. Our five virtual species competed in all pairwise combinations (including with copies of themselves), yielding 15 possible communities. For each of these communities, we simulated the effect of varying strength of stage dependence (11 levels of ***r***_**Δ**_) on each of the five vital rates.

Community dynamics varied greatly depending on the temporal sequence of stochasticity terms, which may obscure the effect of stage dependence. In each simulation, environmental stochasticity was implemented as additive noise on the focal vital rate (Eq. 2), where annual stochasticity terms were drawn independently for each year from a normal distribution with mean zero and standard deviation equal to 10% of the minimum baseline value of that vital rate (Eq. 9). Stochasticity terms were independent across time, species, and replicate subsets, and no temporal autocorrelation was imposed. To ensure that differences in community dynamics across values of ***r***_**Δ**_ were attributable to stage dependence rather than to different stochastic realisations, we generated 10 environmental sequences. Each environmental sequence contains the appropriately-scaled additive stochastic effects (***ϵ***_***θ***,***i***_)on all five vital rates for both species, for 2000 time steps. We then duplicated those 10 sequences, swapping the focal and competitor species, yielding 20 total environmental sequences. We refer to these 20 environmental sequences as “replicates” that are applied to all combinations of two-species communities, density-dependent vital rates, and levels of stage dependence.

To minimise the effect of non-equilibrium initial conditions, we started each simulation from the equilibrium population size (***N*\*** = 500) and the equilibrium structures that were numerically calculated in a preliminary set of simulations (see Model parameterisation, above). To obtain a long-term time-series sample, we sampled species and stage-specific abundances (***N***_**θ**,***i***_**(*t*)**) from ***t*** = 600 to ***t*** = 2,000 time steps (n = 1,400). We then pooled the data across the 20 stochastic environment replicates to provide a large time series dataset (n = 28,000) for each combination of two-species community, density-dependent focal vital rate, and level of stage dependence.

To test our hypothesis H1, we fit community models with or without stage-dependent interactions to each pooled time series and then quantified the predictive accuracy of the fitted model. Specifically, we fit the following linear regression models:

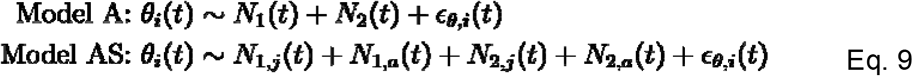

Model A represents a model without stage-dependent interaction, while in Model AS stage-dependent interactions are estimated. After determining the slope estimates, we used each model to predict community dynamics from ***t***_**0**_ = 600 years to ***t***_**end**_ = 700 years within each replicate. To do so, we first set the initial stage-specific abundances to be equal to those at ***t*** = 600 years in the original simulated time-series data. Then, in each year ***t***, we calculated the predicted density-dependent vital rate ***θ*** of each species with the slope and intercept estimates of each model, along with the community structure in that year predicted by the focal model and the true stochasticity term ***ϵ***_***θ***,***i***_**(*t*)** from the original simulation. Finally, with the predicted density-dependent vital rate, we constructed MPMs following Eq. 1 and calculated the predicted community structure at time ***t* + 1**. We repeated this procedure to obtain two predicted time series corresponding to either model A or AS for each original time series.

To quantify the accuracy of each model, we calculated the mean absolute percentage error (MAPE) of the predicted time series relative to the original time series. The MAPE for each species ***i*** was calculated as follows:

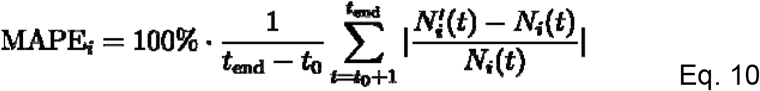

Here, 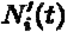 is the abundance of species ***i*** at time ***t*** as predicted by the model and **N**_***i***_**(*t*)** is the abundance of species ***i*** at time ***t*** in the original time series data.

### Analysis software

We conducted all data analyses and community simulations using R (v. 4.4.1). Data manipulation and visualisation were implemented using the *tidyverse* family of R packages (v. 2.0.0) (Wickham et al., 2019). GLMMs were fit and analysed using the *lme4* (v. 1.1.35.5) (Bates et al., 2015) and *car* (v. 3.1.3) (Fox & Weisberg, 2019) packages. Demographic variables were calculated from MPMs using the *popbio* (v. 2.8) (Stubben & Milligan, 2007) and *Rage* (v. 1.6.0) (Jones et al., 2022) R packages. We carried out algebraic expansions, solved equations, and evaluated derivatives and integrals using Wolfram Mathematica (v. 14.2.0.0).

## Results

We quantified how stage dependence in biotic interactions and life history strategy influence the predictive accuracy of community models with or without stage-dependent interactions. Across all simulations, both models closely reproduced the stochastic community dynamics, but prediction error varied systematically with the strength of stage dependence and, in some cases, with the life history of the focal species (Fig. 3). Additionally, we observed that temporal variation in population structure was relatively small (coefficient of variation < 0.036) but variable depending on the life history of the focal species (Fig. 4).

**Figure 3.**
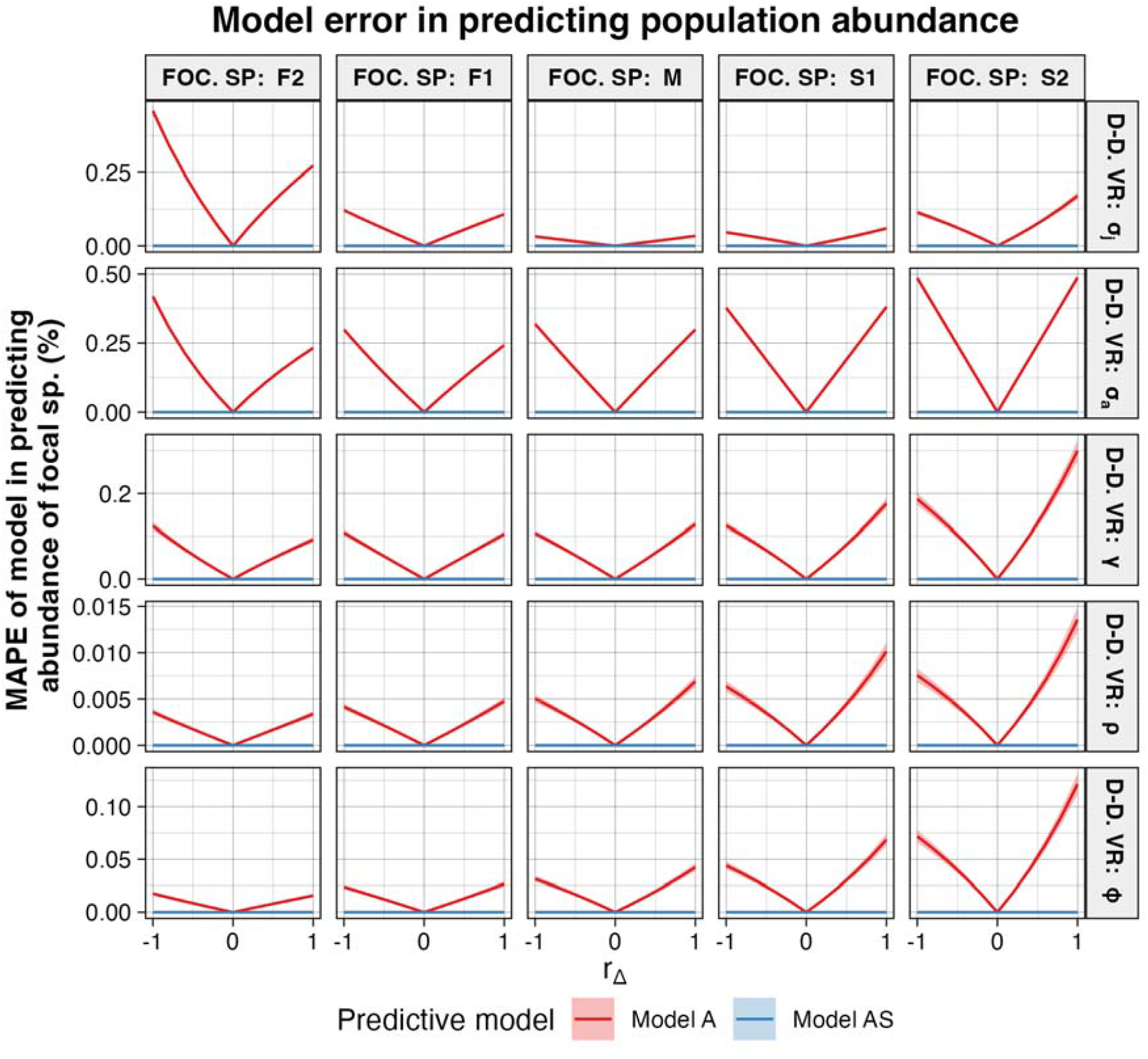
As expected by our hypothesis H1, the predictive error of a community model without stage-dependent interactions (Model A) increases with the strength of stage dependence, but the effect of stage dependence varies with the life history of the focal species and the density-dependent vital rate. To quantify the impact of stage-dependent interactions on stochastic community dynamics, we fit community projection models with (Model AS) or without (Model A) stage-dependent interactions and quantified their accuracy in predicting a 100-year simulated time series. The x-axis represents stage dependence, including simulations where only adults exert density-dependence (***r***_**Δ**_ = −1), where juveniles and adults exert equal density-dependence (***r***_**Δ**_ = 0), and where only juveniles exert density-dependence (***r***_**Δ**_ = 1). The y-axis represents the mean absolute percentage error (MAPE) of each predictive model in predicting the original time series data. Columns represent the life history of the focal species whose abundance is being modelled, varying between the fastest life history (F2) to the slowest life history (S2). Since the life history of the non-focal competitor species did not impact the two models’ predictive accuracy (Fig. S2), the data have been pooled across different competitor identities. Rows represent the density-dependent vital rate of juvenile (***σ***_***j***_) and adult (***σ***_***a***_) survival, progression (***γ***), retrogression (***ρ***), and fertility (***ϕ***). The solid lines indicate the mean MAPE across replicates, and the 95% C.I. is shown as the shaded area.

**Figure 4.**
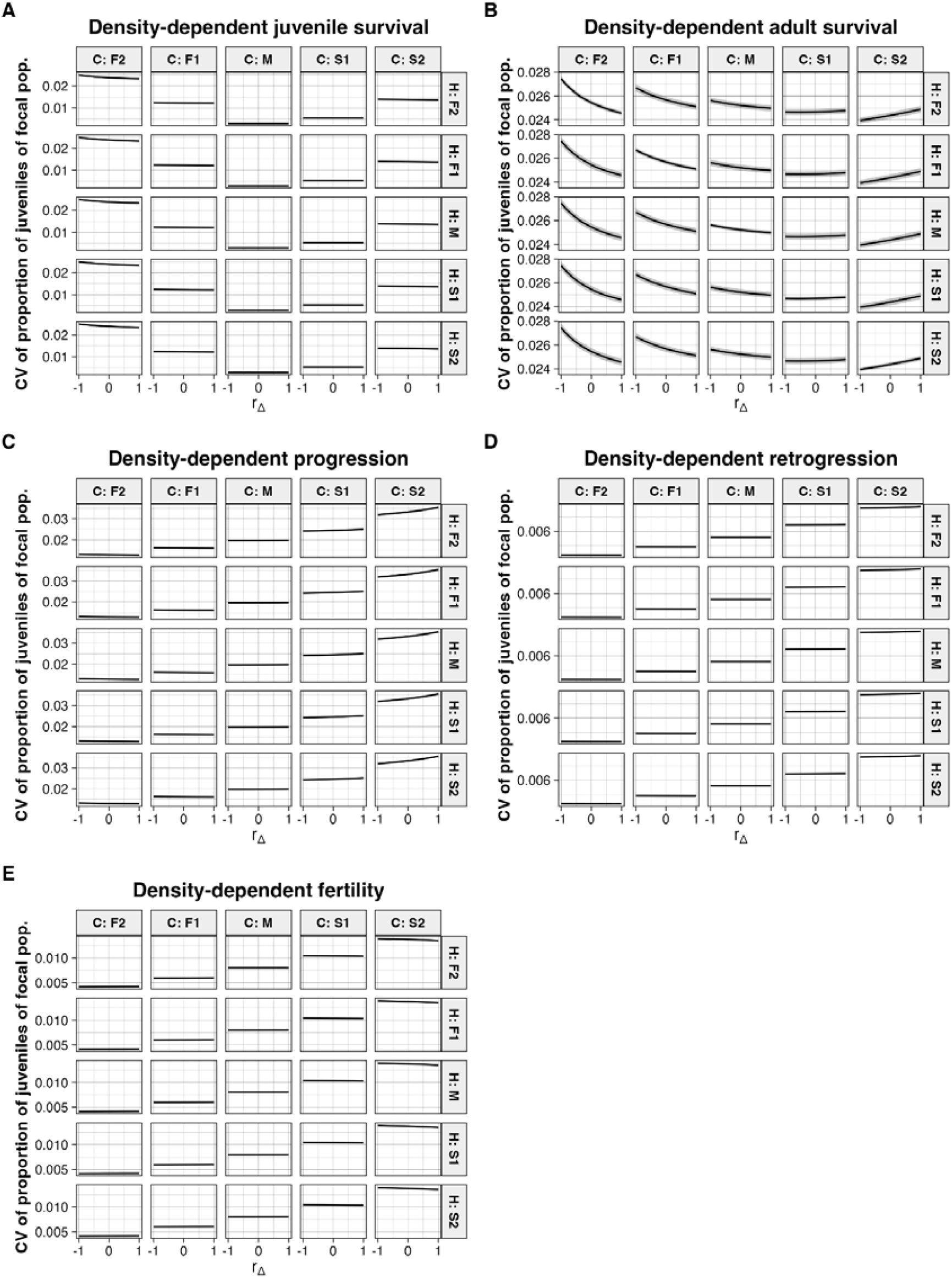
Under our simulations, temporal variation in population structures around respective equilibria was small but variable in a life history-dependent manner. This figure shows the coefficient of variation (CV) in the structure of the focal population across all possible combinations of life histories and density-dependent vital rates. The x-axis represents stage dependence, including simulations where only adults exert density-dependence (***r***_**Δ**_ = −1), where juveniles and adults exert equal density-dependence (***r***_**Δ**_ = 0), and where only juveniles exert density-dependence (***r***_**Δ**_ = 1). The y-axis represents the coefficient of variation in the structure of the focal population as quantified by the proportion of juveniles across the whole time frame of each simulation. The columns denote the life history of the focal population from fastest (F2) to slowest (S2), and the rows denote the life history of the competing population from fastest (F2) to slowest (S2). These figures show the temporal variation in population structure for density-dependent (**A**) juvenile (***σ***_***j***_) and (**B**) adult (***σ***_***a***_) survival, (**C**) progression (***γ***), (**D**) retrogression (***ρ***), and (**E**) fertility (***ϕ***). The solid lines represent the mean coefficient of variation across replicates, and the 95% C.I. is shown as the shaded area.

### Hypothesis 1 (H1): effect of stage dependence on predictive accuracy

Consistent with H1, increasing the strength of stage dependence (***r***_**Δ**_) led to a monotonic increase in the mean absolute percentage error (MAPE) of the community model without stage-dependent interactions (Model A) across all density-dependent vital rates (Fig. 3, 5). When juveniles and adults exerted equal per-capita density-dependent effects (***r***_**Δ**_ = 0), Model A performed nearly as well as the model with stage dependence (Model AS). However, as interactions became increasingly stage dependent, with density dependence driven primarily by either juveniles (***r***_**Δ**_ = 1) or adults (***r***_**Δ**_ = −1), the predictive accuracy of Model A declined.

**Figure 5.**
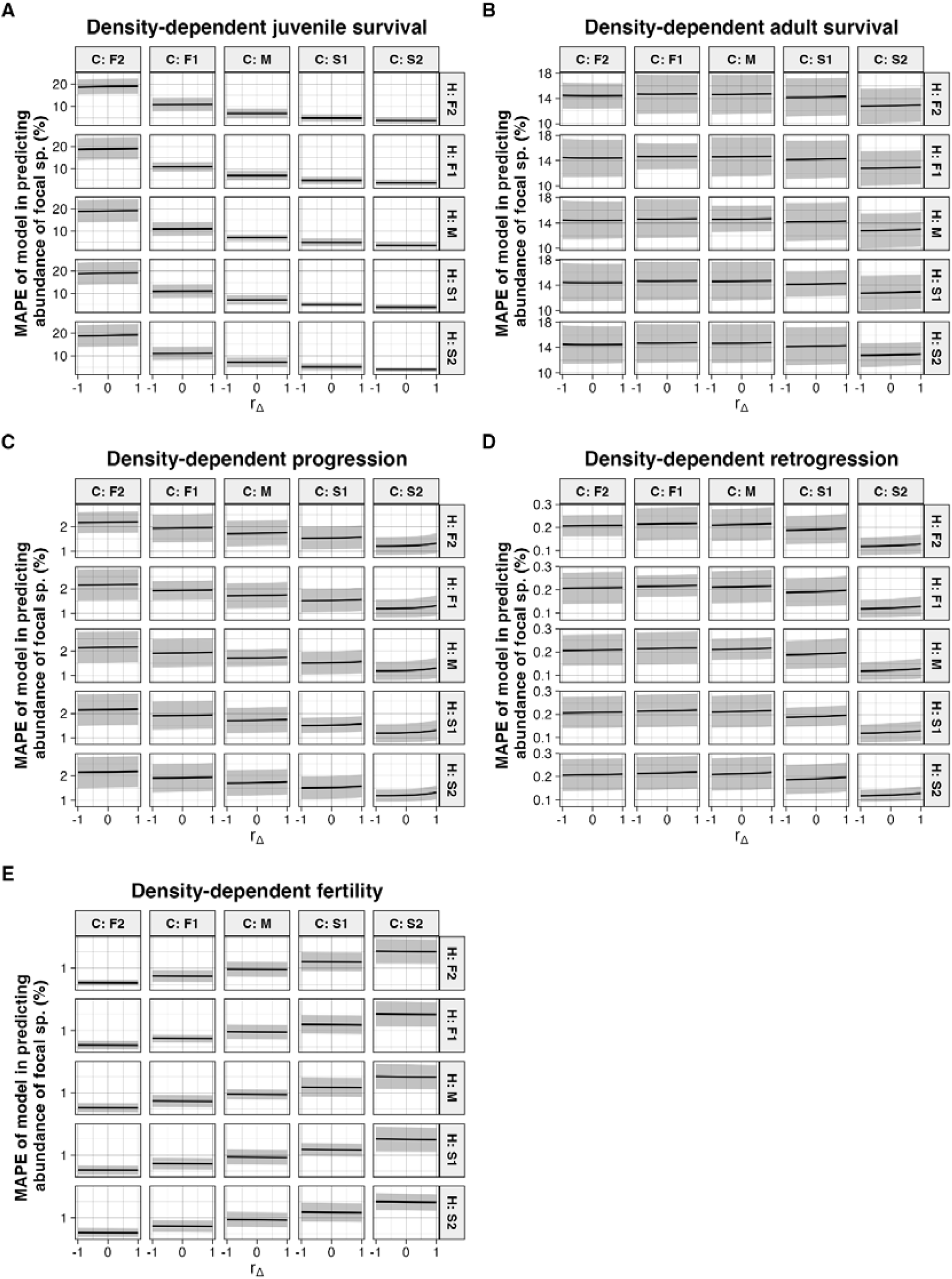
Stage-dependent interactions systematically increase forecasting error in simplified community models, but the magnitude of this error remains small across life histories and demographic pathways. Mean absolute percentage error (MAPE) of the community model without stage-dependent interactions (Model A) as a function of the strength of stage dependence (***r***_**Δ**_), shown separately for five alternative density-dependent vital rates: juvenile survival, adult survival, progression, retrogression, and fertility. The parameter determines the relative per-capita competitive effect of juveniles versus adults, ranging from adult-dominated interactions (***r***_**Δ**_ = −1) to equal effects across stages (***r***_**Δ**_ = 0) to juvenile-dominated interactions (***r***_**Δ**_ = 1). Lines represent different life-history strategies of the focal species; shading (if applicable) indicates variation across stochastic replicates. Across all vital-rate pathways, prediction error increases monotonically with the magnitude of stage asymmetry (|***r***_**Δ**_|). However, absolute error remains below 0.7% in all simulations. Differences among life-history strategies depend on which vital rate mediates density dependence, with contrasting sensitivities observed for juvenile survival versus progression, retrogression, and fertility. The stage-structured model including stage dependence (Model AS) achieved near-zero error in all scenarios (not shown).

Across all vital-rate pathways, the increase in prediction error was smooth and approximately symmetric around ***r***_**Δ**_ = 0, indicating that forecasting sensitivity depended on the magnitude of stage asymmetry rather than on whether juveniles or adults exerted stronger competitive effects. Model AS consistently achieved near-zero error (MAPE < 0.0003%), reflecting the close correspondence between its structure and the data-generating process, whereas the MAPE of Model A remained below 0.7% even under maximal stage dependence. Thus, although ignoring stage dependence introduced systematic forecasting error, its quantitative impact on long-term stochastic projections was limited.

The rate at which MAPE increased with |***r***_**Δ**_ | varied among density-dependent vital rates (Fig. 3, 5). When density dependence operated through juvenile survival, prediction error rose more steeply with increasing stage asymmetry than when it operated through adult survival. Progression, retrogression, and fertility produced intermediate but distinct patterns, indicating that the demographic pathway through which competition acts mediates how strongly stage asymmetry translates into forecasting divergence. Nonetheless, even in the most sensitive scenarios, absolute errors remained small relative to total population size.

### Hypothesis 2 (H2): interaction between stage dependence and life history strategy

Contrary to H2, faster life histories did not universally exacerbate the effect of stage dependence on the predictive accuracy of Model A. Instead, the relationship between life history and prediction error depended on which vital rate was density dependent (Fig. 3).

When juvenile survival was density dependent, faster life histories of the focal species were generally associated with larger MAPE values at a given level of stage dependence. In these cases, concentrating competitive effects on a stage that strongly contributes to short-term recruitment amplified discrepancies when stage dependence was ignored. As ***r***_**Δ**_ moved away from zero, the divergence between life histories became more pronounced, with faster strategies showing steeper increases in prediction error.

In contrast, when progression, retrogression, or fertility was density dependent, slower life histories tended to exhibit greater prediction error. Under these pathways, misrepresenting the stage distribution of competitive effects propagated more strongly through demographic strategies that rely on adult persistence and stage transitions for population growth. Consequently, slower life histories showed greater sensitivity to increasing |***r***_**Δ**_ | than faster ones.

When adult survival was density dependent, no consistent relationship emerged between life history pace and predictive accuracy, and differences among life histories were comparatively small across the full range of ***r***_**Δ**_.

In all cases, the life history strategy of the non-focal competitor species had negligible effects on the predictive accuracy of both models (Fig. S2). Differences in forecasting performance were therefore primarily driven by the demographic characteristics of the focal species and by the vital rate through which density dependence was imposed.

## Discussion

Biotic interactions are a central driver of community dynamics and coexistence, and their strength and direction often depend on the ontogenetic stage of interacting individuals (Miller & Rudolf, 2011; Nakazawa, 2015). Understanding when such stage-dependent interactions meaningfully influence community dynamics under environmental stochasticity is therefore critical for determining the level of demographic detail required for accurate ecological forecasting (Petchey et al., 2015; Getz et al., 2018). As such, here we quantified the impact of stage dependence on the importance of stage-specific interactions in predicting stochastic community dynamics. We address a current knowledge gap in the field by modelling the joint effect of stage-dependence and stochasticity on community dynamics. We extend density-dependent (Jensen, 1995; Miller et al., 2002; Neubert & Caswell, 2000; Smouse & Weiss, 1975) and stochastic (Fieberg & Ellner, 2001; Nakaoka, 1996; Sykes, 1969) matrix population models (MPMs) to determine how stage dependence influences the predictive accuracy of community models with or without stage-dependent biotic interactions. We show that stronger stage-dependence can increase the error of a community model without stage-specific biotic interactions, although this error is proportionally marginal, but more important in slow-living species. Together, the results suggest that, although stage dependence can impact stochastic community dynamics and thereby the accuracy of predictive models, community models without stage-dependent interactions may be adequate for ecological forecasts.

The impact of stage dependence may be best explained by the amount of temporal variation in population structure. We found that the error of a community model without stage-dependent interactions (Model A) was not always greater when the focal species had a faster life history. The difference in predictive error of model A across focal species was better explained by the amount of temporal variation in population structure. That is, focal species with greater temporal variation in population structure were associated with greater error of model A (Fig. 4, S3). This pattern is consistent with the logical basis of our hypothesis H2, that greater deviation of population structure from its equilibrium increases the discrepancy between the true per-capita average interaction strength and the corresponding estimate without stage-dependent interactions.

In turn, the amount of temporal variation in population structure may be impacted by intrinsic features of each species’ life history. Notably, the inverse relationship between the pace of life and variation in population structure seen with density-dependent progression (***γ***), retrogression (***ρ***), and fertility (***ϕ***) contrasts with previous empirical results (Bjørkvoll et al., 2012; Sæther et al., 2004; Sæther et al., 2013). In our model, temporal variation in population structure may be influenced by the sensitivity of the stable population structure (*sensu* Caswell 2001) to changes in the density-dependent vital rate. Here, we derived this quantity by differentiating an expression for the proportion of juveniles at stable population structure in terms of each of the five vital rates (Supplementary information S3). For example, when progression (***γ***), retrogression (***ρ***) or fertility (***ϕ***) were density-dependent, the sensitivity of the stable population structure at the realised mean vital rates was 5-to 94-fold greater in the species with the slowest life history compared to the fastest life history (Table S2). Since these virtual life histories are not reflective of real species, the generality of this pattern in natural communities cannot be verified. To address this question, the present work may be repeated using empirical vital rates that reflect ecological and evolutionary constraints on life history strategy (Bielby et al., 2007; Salguero-Gómez et al., 2016). Moreover, it remains unclear whether insights derived from two-species systems transfer to more diverse communities, where indirect interactions, higher-order effects, and network structure may amplify or dampen the influence of stage-dependent density dependence. Generally, we suggest that quantifying vital rate-specific density-dependent effects and metrics such as the sensitivity of stable population structure may reveal such complex relationships between life history and stochastic community dynamics.

The low predictive error of the community model without stage-dependent interactions is likely driven by the large influence of environmental stochasticity and the limited variation in population structure observed in our simulations. Across all scenarios, the predictive error remained extremely small (< 0.7% MAPE), even under strong stage-dependence in the data-generating process. Such marginal error suggests that stage dependence did not substantially alter stochastic community dynamics under the simulated conditions. This result contrasts with previous theoretical works demonstrating that stage-dependent interactions can qualitatively reshape community dynamics, including stability, coexistence conditions, and alternative stable states (de Roos, 2021; Loreau & Ebenhoh, 1994; Moll & Brown, 2008; Nakazawa, 2011a, 2011b; Schellekens et al., 2010; Wollrab et al., 2013). In our model, stochastic fluctuations in vital rates appeared to dominate dynamics, while variation in stage structure remained small (coefficient of variation < 0.036), limiting the extent to which stage-dependent effects could translate into changes in population-level density dependence.

Importantly, our simulations were initialised at stationary equilibrium abundances and stage structures. Consequently, our conclusions apply to communities fluctuating around demographic equilibria rather than to systems undergoing invasion, recovery, or strong transient dynamics. The ability of species to establish from low density, recover following disturbance, or exhibit transient amplification may itself depend on the interaction between stochasticity and stage-dependent density dependence. Under such non-equilibrium conditions, demographic structure could exert stronger effects on competitive outcomes and forecast accuracy than observed here. Exploring how stage dependence influences invasion probability, exclusion dynamics, or transient trajectories represents an important direction for future work.

In conservation, stage-dependent interactions may be more important in predicting stochastic community dynamics when anthropogenic drivers cause non-equilibrium population structures. Examples of such drivers with stage-dependent intensity include hunting (Torres-Porras et al., 2014), selective logging (Tsingalia, 2010), and fishing (Hsieh et al., 2010). To investigate whether the performance of the predictive models is influenced by such drivers, future work may quantify predictive accuracy following demographic disturbance. In this context, metrics of transient population dynamics (*sensu* Stott et al. (2011)) may help explain the drivers of variability in population structure across life histories. However, our results suggest that, when there is little fluctuation in population structure, quantifying stage-dependent interactions may not improve the accuracy of community forecasts.

In conclusion, stage-dependence in biotic interactions can influence stochastic community dynamics and their predictive modelling, although its quantitative importance depends on the magnitude of structural fluctuations. The present empirical and theoretical methodologies contribute to a growing body of literature integrating demography and community ecology (de Roos, 2021; Lytle & Tonkin, 2023; Miller & Rudolf, 2011). In this study, we parameterised the system such that both species persisted at a non-zero equilibrium, thereby focusing on forecast accuracy conditional on coexistence. However, the same modelling framework could be used to ask a complementary question: how does stage dependence alter invasion growth rates and the probability of coexistence under stochastic dynamics? Extending this approach to evaluate whether stage-dependent interactions expand or contract coexistence regions would provide a direct link to Modern Coexistence Theory (Chesson, 2000, 2003; Ellner et al., 2019). More broadly, approaches similar to the current model may help bridge stochastic coexistence theory with life-history theory, including the fast–slow continuum (Gaillard et al., 1989; Salguero-Gómez et al., 2016) and emerging perspectives on demographic diversity beyond simple pace-of-life axes (Stott et al., 2024; Van de Walle et al., 2023).

## Supporting information

SOMs

## Acknowledgments

We thank S. Ellner, P. Adler, R. Snyder, G. Hooker, and G. Barabás for their feedback on the theoretical basis of this work. RSG and CMH were supported by a NERC Pushing the Frontiers grant (NE/X013766/1) to RSG. CMH was also partially supported by a Marie Curie Fellowship (MSCA DensPopDy #10115386) with funding through UKRI (EP/Z002826/1) hosted by RSG. CAR was supported by the US National Science Foundation (#2410511). CAR and BWB were supported by a LTREB grant from the U.S. National Science Foundation (DEB□2425575).

