## Supplementary material for "Stage-dependent biotic interactions may not be important for stochastic competitive dynamics with little variation in stage structure": SOMs

### Supplementary information S1. Sensitivities of the asymptotic population growth rate <https://www.codecogs.com/eqnedit.php?latex=%5Clambda#0> to each vital rate

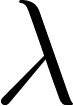

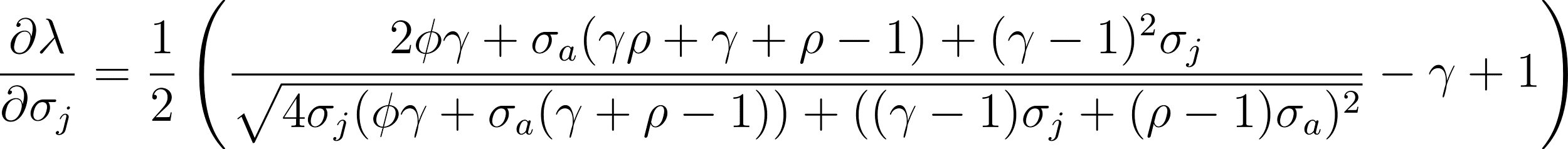
<https://www.codecogs.com/eqnedit.php?latex=%5C%5C%5C%5C%20%5Cfrac%7B%5Cpartial%20%5Clambda%7D%7B%5Cpartial%20%5Csigma_j%7D%20%3D%20%5Cfrac%7B1%7D%7B2%7D%20%5Cleft(%5Cfrac%7B2%20%5Cphi%20%5Cgamma%2B%5Csigma_a%20(%5Cgamma%20%5Crho%2B%5Cgamma%2B%5Crho-1)%2B(%5Cgamma-1)%5E2%20%5Csigma_j%7D%7B%5Csqrt%7B4%20%5Csigma_j%20(%5Cphi%20%5Cgamma%2B%5Csigma_a%20(%5Cgamma%2B%5Crho-1))%2B((%5Cgamma-1)%20%5Csigma_j%2B(%5Crho-1)%20%5Csigma_a)%5E2%7D%7D-%5Cgamma%2B1%5Cright)%5C%5C%5C%5C%20#0>

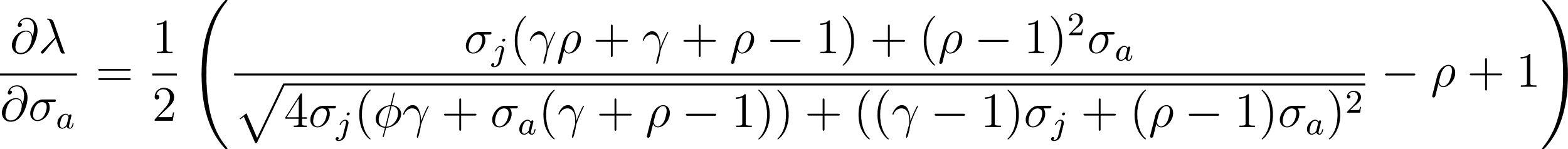
<https://www.codecogs.com/eqnedit.php?latex=%5C%5C%5C%5C%20%5Cfrac%7B%5Cpartial%20%5Clambda%7D%7B%5Cpartial%20%5Csigma_a%7D%20%3D%20%5Cfrac%7B1%7D%7B2%7D%20%5Cleft(%5Cfrac%7B%5Csigma_j%20(%5Cgamma%20%5Crho%2B%5Cgamma%2B%5Crho-1)%2B(%5Crho-1)%5E2%20%5Csigma_a%7D%7B%5Csqrt%7B4%20%5Csigma_j%20(%5Cphi%20%5Cgamma%2B%5Csigma_a%20(%5Cgamma%2B%5Crho-1))%2B((%5Cgamma-1)%20%5Csigma_j%2B(%5Crho-1)%20%5Csigma_a)%5E2%7D%7D-%5Crho%2B1%5Cright)%5C%5C%5C%5C%20#0>

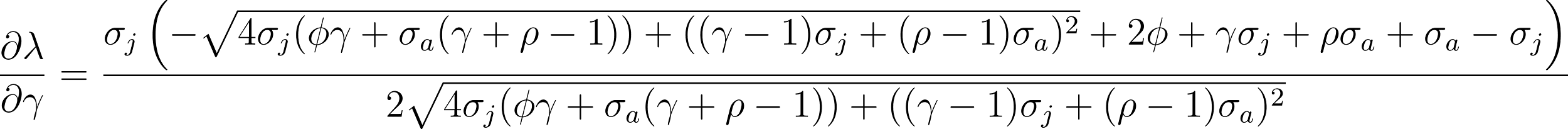
<https://www.codecogs.com/eqnedit.php?latex=%5C%5C%5C%5C%20%5Cfrac%7B%5Cpartial%20%5Clambda%7D%7B%5Cpartial%20%5Cgamma%7D%20%3D%20%5Cfrac%7B%5Csigma_j%20%5Cleft(-%5Csqrt%7B4%20%5Csigma_j%20(%5Cphi%20%5Cgamma%2B%5Csigma_a%20(%5Cgamma%2B%5Crho-1))%2B((%5Cgamma-1)%20%5Csigma_j%2B(%5Crho-1)%20%5Csigma_a)%5E2%7D%2B2%20%5Cphi%2B%5Cgamma%20%5Csigma_j%2B%5Crho%20%5Csigma_a%2B%5Csigma_a-%5Csigma_j%5Cright)%7D%7B2%20%5Csqrt%7B4%20%5Csigma_j%20(%5Cphi%20%5Cgamma%2B%5Csigma_a%20(%5Cgamma%2B%5Crho-1))%2B((%5Cgamma-1)%20%5Csigma_j%2B(%5Crho-1)%20%5Csigma_a)%5E2%7D%7D%5C%5C%5C%5C%20#0>

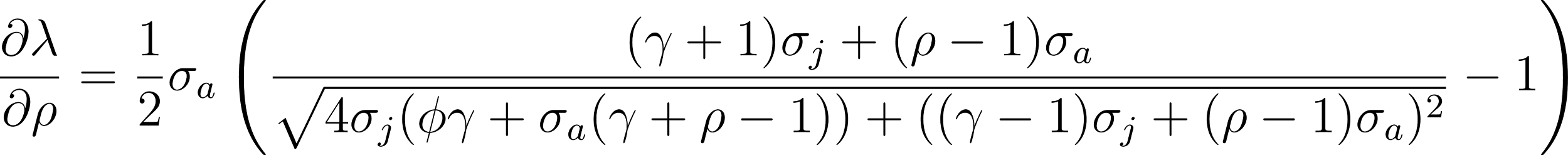
<https://www.codecogs.com/eqnedit.php?latex=%5C%5C%5C%5C%20%5Cfrac%7B%5Cpartial%20%5Clambda%7D%7B%5Cpartial%20%5Crho%7D%20%3D%20%5Cfrac%7B1%7D%7B2%7D%20%5Csigma_a%20%5Cleft(%5Cfrac%7B(%5Cgamma%2B1)%20%5Csigma_j%2B(%5Crho-1)%20%5Csigma_a%7D%7B%5Csqrt%7B4%20%5Csigma_j%20(%5Cphi%20%5Cgamma%2B%5Csigma_a%20(%5Cgamma%2B%5Crho-1))%2B((%5Cgamma-1)%20%5Csigma_j%2B(%5Crho-1)%20%5Csigma_a)%5E2%7D%7D-1%5Cright)%5C%5C%5C%5C%20#0>

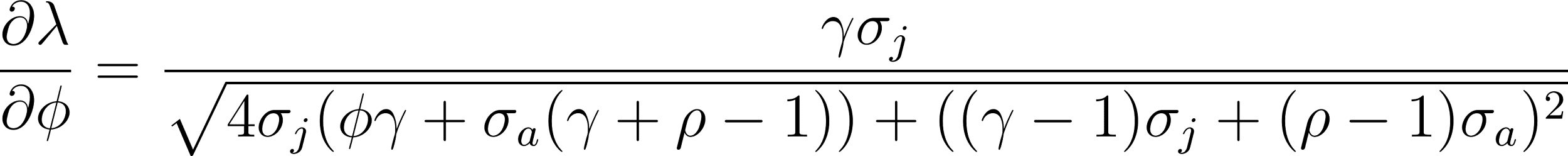
<https://www.codecogs.com/eqnedit.php?latex=%5C%5C%5C%5C%20%5Cfrac%7B%5Cpartial%20%5Clambda%7D%7B%5Cpartial%20%5Cphi%7D%20%3D%20%5Cfrac%7B%5Cgamma%20%5Csigma_j%7D%7B%5Csqrt%7B4%20%5Csigma_j%20(%5Cphi%20%5Cgamma%2B%5Csigma_a%20(%5Cgamma%2B%5Crho-1))%2B((%5Cgamma-1)%20%5Csigma_j%2B(%5Crho-1)%20%5Csigma_a)%5E2%7D%7D%5C%5C%5C%5C%20#0>

### **Table S1. Baseline interaction coefficients define the strength of density dependence prior to imposing stage asymmetry.** Baseline per-capita interaction coefficients used to parameterise density dependence for each focal vital rate (juvenile survival (<https://www.codecogs.com/eqnedit.php?latex=%5Csigma_j#0>), adult survival (<https://www.codecogs.com/eqnedit.php?latex=%5Csigma_a#0>), progression (<https://www.codecogs.com/eqnedit.php?latex=%5Cgamma#0>), retrogression (<https://www.codecogs.com/eqnedit.php?latex=%5Crho#0>), and fertility (<https://www.codecogs.com/eqnedit.php?latex=%5Cphi#0>)). These coefficients determine the magnitude of intra- and interspecific competitive effects under the reference condition in which juveniles and adults exert equal per-capita effects (<https://www.codecogs.com/eqnedit.php?latex=r_%5CDelta#0> = 0). Values are specified for each species and density-dependent pathway and serve as the starting point for generating stage-dependent interactions through the stage-weighting parameter (<https://www.codecogs.com/eqnedit.php?latex=r_%5CDelta#0>). All subsequent simulations modify the relative contributions of juveniles and adults while preserving the overall interaction strength defined by these baseline coefficients.

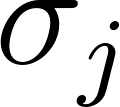

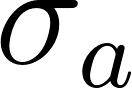

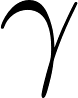

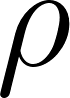

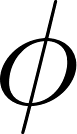

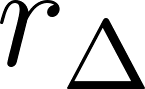

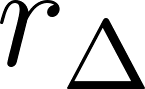

| D-D  vital rate | Species | 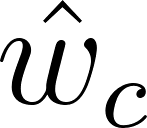<https://www.codecogs.com/eqnedit.php?latex=%5Chat%7Bw%7D_c#0> | Peak 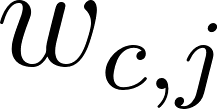<https://www.codecogs.com/eqnedit.php?latex=w_%7Bc%2Cj%7D#0> | Peak 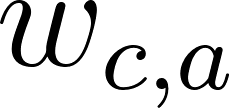<https://www.codecogs.com/eqnedit.php?latex=w_%7Bc%2Ca%7D#0> | 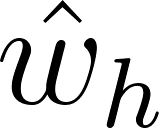<https://www.codecogs.com/eqnedit.php?latex=%5Chat%7Bw%7D_h#0> | Peak 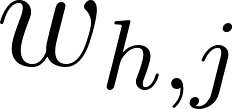<https://www.codecogs.com/eqnedit.php?latex=w_%7Bh%2Cj%7D#0> | Peak 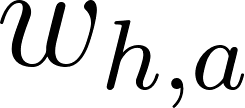<https://www.codecogs.com/eqnedit.php?latex=w_%7Bh%2Ca%7D#0> |
| --- | --- | --- | --- | --- | --- | --- | --- |
| 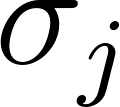<https://www.codecogs.com/eqnedit.php?latex=%5Csigma_j#0> | F2 | -1.96e-04 | -3.11e-04 | -5.31e-04 | -4.36e-05 | -9.22e-05 | -8.82e-05 |
| 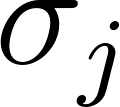<https://www.codecogs.com/eqnedit.php?latex=%5Csigma_j#0> | F1 | -1.96e-04 | -3.64e-04 | -4.26e-04 | -4.36e-05 | -9.79e-05 | -8.08e-05 |
| 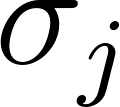<https://www.codecogs.com/eqnedit.php?latex=%5Csigma_j#0> | M | -1.96e-04 | -4.17e-04 | -3.71e-04 | -4.36e-05 | -1.04e-04 | -7.61e-05 |
| 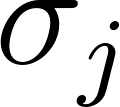<https://www.codecogs.com/eqnedit.php?latex=%5Csigma_j#0> | S1 | -1.96e-04 | -4.68e-04 | -3.38e-04 | -4.36e-05 | -1.09e-04 | -7.29e-05 |
| 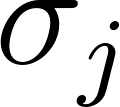<https://www.codecogs.com/eqnedit.php?latex=%5Csigma_j#0> | S2 | -1.96e-04 | -5.13e-04 | -3.18e-04 | -4.36e-05 | -1.14e-04 | -7.07e-05 |
| 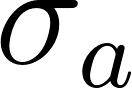<https://www.codecogs.com/eqnedit.php?latex=%5Csigma_a#0> | F2 | -2.95e-04 | -4.71e-04 | -7.86e-04 | -6.55e-05 | -1.18e-04 | -1.49e-04 |
| 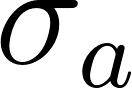<https://www.codecogs.com/eqnedit.php?latex=%5Csigma_a#0> | F1 | -2.95e-04 | -5.16e-04 | -6.87e-04 | -6.55e-05 | -1.21e-04 | -1.42e-04 |
| 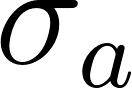<https://www.codecogs.com/eqnedit.php?latex=%5Csigma_a#0> | M | -2.95e-04 | -5.50e-04 | -6.34e-04 | -6.55e-05 | -1.24e-04 | -1.39e-04 |
| 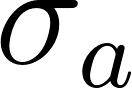<https://www.codecogs.com/eqnedit.php?latex=%5Csigma_a#0> | S1 | -2.95e-04 | -5.67e-04 | -6.13e-04 | -6.55e-05 | -1.24e-04 | -1.38e-04 |
| 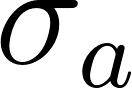<https://www.codecogs.com/eqnedit.php?latex=%5Csigma_a#0> | S2 | -2.95e-04 | -5.54e-04 | -6.29e-04 | -6.55e-05 | -1.23e-04 | -1.40e-04 |
| 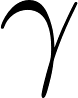<https://www.codecogs.com/eqnedit.php?latex=%5Cgamma#0> | F2 | -1.47e-04 | -2.58e-04 | -3.43e-04 | -3.27e-05 | -6.88e-05 | -6.37e-05 |
| 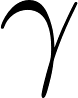<https://www.codecogs.com/eqnedit.php?latex=%5Cgamma#0> | F1 | -1.47e-04 | -2.91e-04 | -2.98e-04 | -3.27e-05 | -7.17e-05 | -6.06e-05 |
| 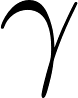<https://www.codecogs.com/eqnedit.php?latex=%5Cgamma#0> | M | -1.47e-04 | -3.20e-04 | -2.73e-04 | -3.27e-05 | -7.40e-05 | -5.87e-05 |
| 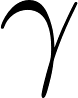<https://www.codecogs.com/eqnedit.php?latex=%5Cgamma#0> | S1 | -1.47e-04 | -3.40e-04 | -2.60e-04 | -3.27e-05 | -7.55e-05 | -5.78e-05 |
| 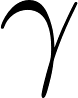<https://www.codecogs.com/eqnedit.php?latex=%5Cgamma#0> | S2 | -1.47e-04 | -3.39e-04 | -2.60e-04 | -3.27e-05 | -7.54e-05 | -5.78e-05 |
| 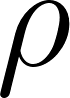<https://www.codecogs.com/eqnedit.php?latex=%5Crho#0> | F2 | 2.45e-05 | 4.47e-05 | 5.45e-05 | 5.45e-06 | 1.25e-05 | 9.97e-06 |
| 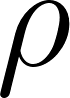<https://www.codecogs.com/eqnedit.php?latex=%5Crho#0> | F1 | 2.45e-05 | 5.10e-05 | 4.73e-05 | 5.45e-06 | 1.31e-05 | 9.44e-06 |
| 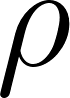<https://www.codecogs.com/eqnedit.php?latex=%5Crho#0> | M | 2.45e-05 | 5.70e-05 | 4.31e-05 | 5.45e-06 | 1.37e-05 | 9.08e-06 |
| 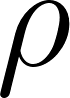<https://www.codecogs.com/eqnedit.php?latex=%5Crho#0> | S1 | 2.45e-05 | 6.24e-05 | 4.04e-05 | 5.45e-06 | 1.43e-05 | 8.83e-06 |
| 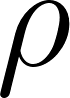<https://www.codecogs.com/eqnedit.php?latex=%5Crho#0> | S2 | 2.45e-05 | 6.62e-05 | 3.90e-05 | 5.45e-06 | 1.47e-05 | 8.67e-06 |
| 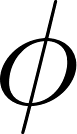<https://www.codecogs.com/eqnedit.php?latex=%5Cphi#0> | F2 | -4.91e-05 | -9.02e-05 | -1.08e-04 | -1.09e-05 | -2.58e-05 | -1.96e-05 |
| 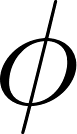<https://www.codecogs.com/eqnedit.php?latex=%5Cphi#0> | F1 | -4.91e-05 | -1.03e-04 | -9.34e-05 | -1.09e-05 | -2.72e-05 | -1.85e-05 |
| 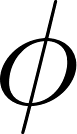<https://www.codecogs.com/eqnedit.php?latex=%5Cphi#0> | M | -4.91e-05 | -1.17e-04 | -8.47e-05 | -1.09e-05 | -2.86e-05 | -1.77e-05 |
| 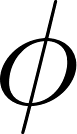<https://www.codecogs.com/eqnedit.php?latex=%5Cphi#0> | S1 | -4.91e-05 | -1.29e-04 | -7.91e-05 | -1.09e-05 | -3.00e-05 | -1.72e-05 |
| 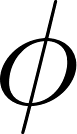<https://www.codecogs.com/eqnedit.php?latex=%5Cphi#0> | S2 | -4.91e-05 | -1.40e-04 | -7.55e-05 | -1.09e-05 | -3.12e-05 | -1.68e-05 |

### Supplementary information S2. Procedure for calculating baseline fertility

When a vital rate other than fertility is density-dependent, we calculated the density-dependent vital rate at the desired equilibrium population size and set
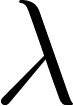
<https://www.codecogs.com/eqnedit.php?latex=%5Clambda#0> = 1, where
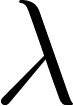
<https://www.codecogs.com/eqnedit.php?latex=%5Clambda#0> is given as:

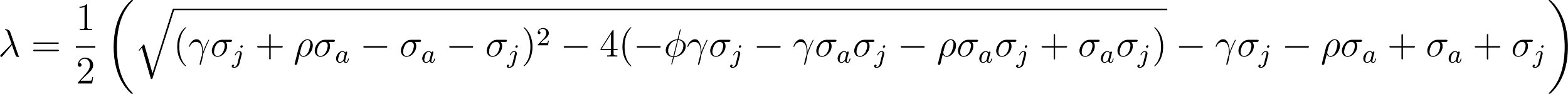
<https://www.codecogs.com/eqnedit.php?latex=%5C%5C%5C%5C%20%5Clambda%20%3D%20%5Cfrac%7B1%7D%7B2%7D%20%5Cleft(%5Csqrt%7B(%5Cgamma%20%5Csigma_j%2B%5Crho%20%5Csigma_a-%5Csigma_a-%5Csigma_j)%5E2-4%20(-%5Cphi%20%5Cgamma%20%5Csigma_j-%5Cgamma%20%5Csigma_a%20%5Csigma_j-%5Crho%20%5Csigma_a%20%5Csigma_j%2B%5Csigma_a%20%5Csigma_j)%7D-%5Cgamma%20%5Csigma_j-%5Crho%20%5Csigma_a%2B%5Csigma_a%2B%5Csigma_j%5Cright)%5C%5C%5C%5C%20#0>. This expression was then solved for fertility (
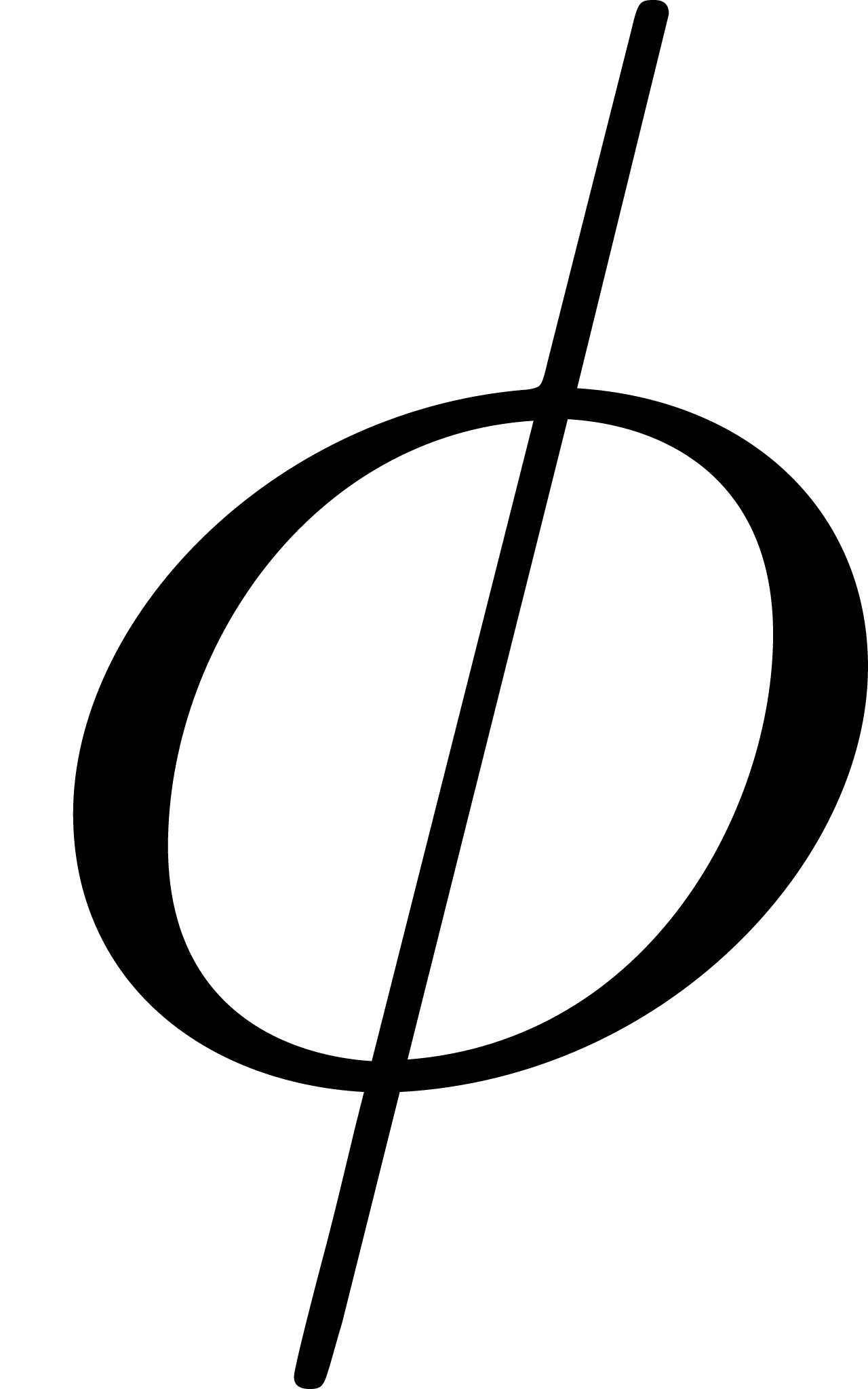
).

When fertility is the density-dependent vital rate, we first calculated the density-dependence term
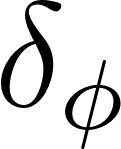
<https://www.codecogs.com/eqnedit.php?latex=%5Cdelta_%5Cphi#0> at the equilibrium population sizes. We replaced
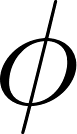
<https://www.codecogs.com/eqnedit.php?latex=%5Cphi#0> above with
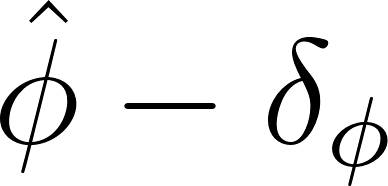
<https://www.codecogs.com/eqnedit.php?latex=%5Chat%7B%5Cphi%7D%20-%20%5Cdelta_%5Cphi#0> and set
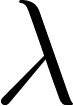
<https://www.codecogs.com/eqnedit.php?latex=%5Clambda#0> = 1, and then solved for
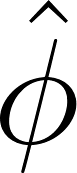
<https://www.codecogs.com/eqnedit.php?latex=%5Chat%7B%5Cphi%7D#0>.

### **Figure S1. Stochastic population dynamics remain stable across life-history combinations and strengths of stage dependence.** Mean total population size of each species across replicate stochastic simulations under varying strengths of stage dependence (<https://www.codecogs.com/eqnedit.php?latex=r_%5CDelta#0>). Each panel corresponds to a specific combination of competitor and focal life-history strategies, denoted as “C: F”, where “C” refers to the competitor species and “F” to the focal species. The numerical suffix (*e.g*., F2, C) identifies the life-history type assigned to that species along the fast-slow continuum defined in the virtual species parameterisation, with faster strategies to the left and slowest to the right. Lines represent mean population size across replicate runs subjected to identical environmental stochasticity sequences, and shaded regions (if shown) indicate variation among replicates. Simulations were initiated at stationary equilibrium and run for 1,400 time steps with environmental stochasticity applied to the density-dependent vital rate. Across all values of <https://www.codecogs.com/eqnedit.php?latex=r_%5CDelta#0> and life-history combinations, populations fluctuated around their equilibrium expectations without sustained directional drift.

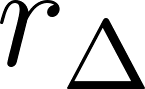

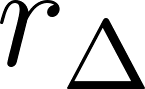

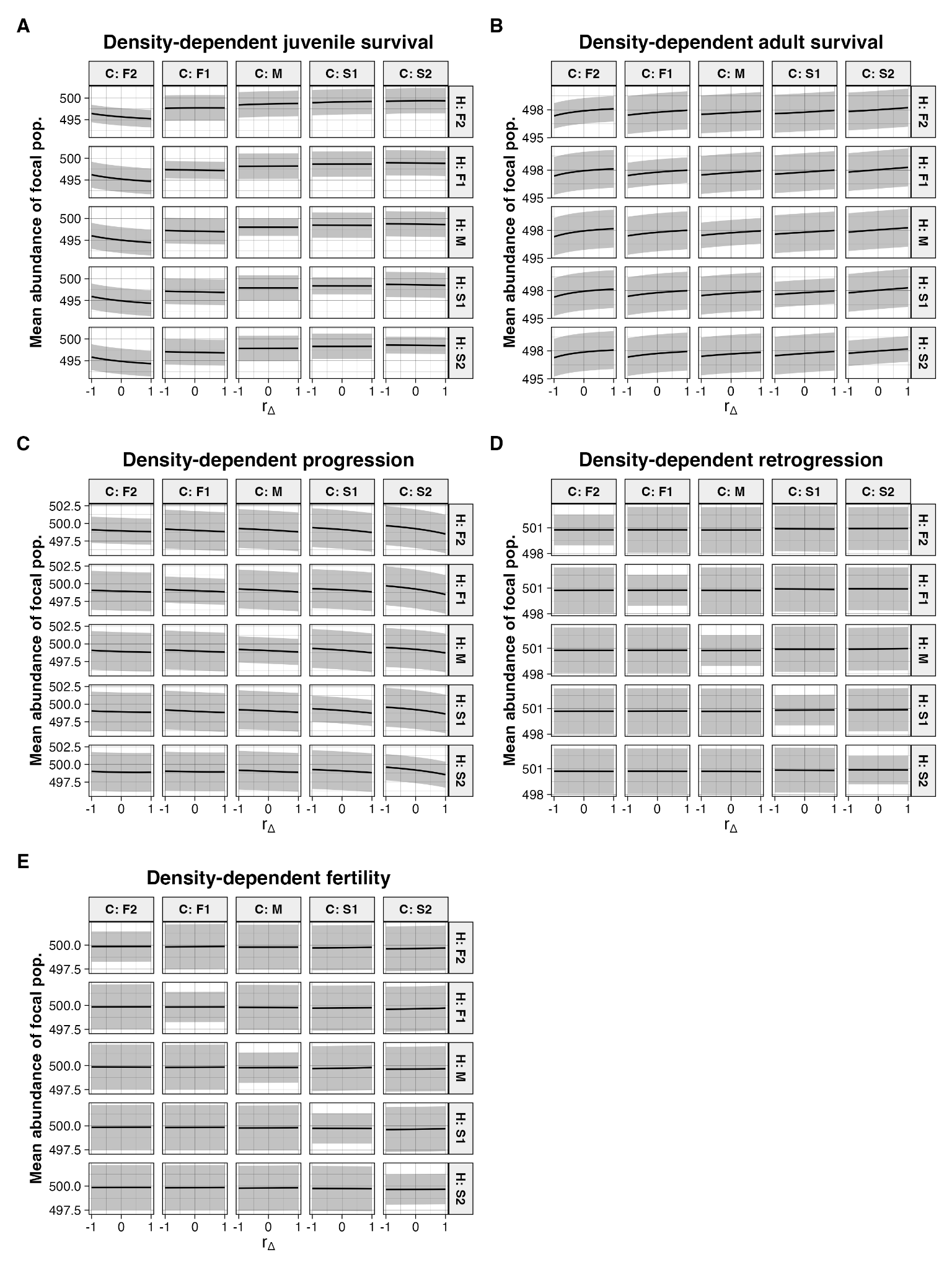

### **Table S2. Life-history strategies differ systematically in their realised demographic properties along the slow–fast continuum.** Mean realised vital rates and associated demographic metrics for each virtual life-history type used in the simulations. Values are calculated from the baseline (density-independent) projection matrices prior to the imposition of stage-dependent competition. Gen. time refers to generation time, calculated as the mean age of reproduction. Life expect. denotes life expectancy. Juv. prop. sens. refers to the proportional sensitivity of population growth rate (λ) to juvenile-stage demographic processes, quantified as the summed elasticities of juvenile survival and progression divided by total elasticity. These metrics characterise the demographic pace and structure of each life-history strategy and define their position along the slow–fast continuum implemented in the virtual species parameterisation.

| D-D vr. | Sp. | 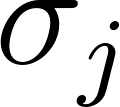<https://www.codecogs.com/eqnedit.php?latex=%5Csigma_j#0> | 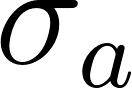<https://www.codecogs.com/eqnedit.php?latex=%5Csigma_a#0> | 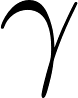<https://www.codecogs.com/eqnedit.php?latex=%5Cgamma#0> | 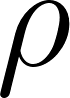<https://www.codecogs.com/eqnedit.php?latex=%5Crho#0> | 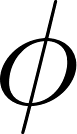<https://www.codecogs.com/eqnedit.php?latex=%5Cphi#0> | Gen. time | Life expect. | Juv. prop. sens. |
| --- | --- | --- | --- | --- | --- | --- | --- | --- | --- |
| 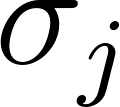<https://www.codecogs.com/eqnedit.php?latex=%5Csigma_j#0> | F2 | 0.28 | 0.60 | 0.90 | 0.05 | 1.63 | 3.35 | 1.66 | -1.33 |
| 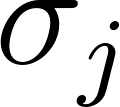<https://www.codecogs.com/eqnedit.php?latex=%5Csigma_j#0> | F1 | 0.41 | 0.69 | 0.75 | 0.06 | 1.01 | 3.92 | 2.15 | -1.23 |
| 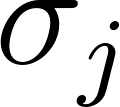<https://www.codecogs.com/eqnedit.php?latex=%5Csigma_j#0> | M | 0.53 | 0.77 | 0.60 | 0.08 | 0.64 | 4.80 | 2.94 | -1.22 |
| 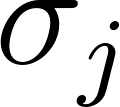<https://www.codecogs.com/eqnedit.php?latex=%5Csigma_j#0> | S1 | 0.66 | 0.86 | 0.45 | 0.09 | 0.39 | 6.26 | 4.46 | -1.31 |
| 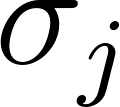<https://www.codecogs.com/eqnedit.php?latex=%5Csigma_j#0> | S2 | 0.78 | 0.95 | 0.30 | 0.10 | 0.19 | 9.10 | 8.70 | -1.58 |
| 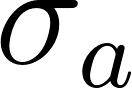<https://www.codecogs.com/eqnedit.php?latex=%5Csigma_a#0> | F2 | 0.40 | 0.42 | 0.90 | 0.05 | 1.58 | 2.71 | 1.69 | 0.23 |
| 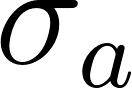<https://www.codecogs.com/eqnedit.php?latex=%5Csigma_a#0> | F1 | 0.53 | 0.51 | 0.75 | 0.06 | 1.12 | 3.06 | 2.07 | 0.28 |
| 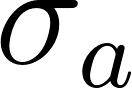<https://www.codecogs.com/eqnedit.php?latex=%5Csigma_a#0> | M | 0.65 | 0.60 | 0.60 | 0.08 | 0.81 | 3.58 | 2.67 | 0.33 |
| 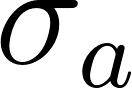<https://www.codecogs.com/eqnedit.php?latex=%5Csigma_a#0> | S1 | 0.78 | 0.68 | 0.45 | 0.09 | 0.56 | 4.40 | 3.72 | 0.39 |
| 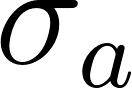<https://www.codecogs.com/eqnedit.php?latex=%5Csigma_a#0> | S2 | 0.90 | 0.77 | 0.30 | 0.10 | 0.34 | 5.97 | 6.23 | 0.43 |
| 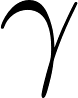<https://www.codecogs.com/eqnedit.php?latex=%5Cgamma#0> | F2 | 0.40 | 0.60 | 0.81 | 0.05 | 1.20 | 3.41 | 1.95 | 0.13 |
| 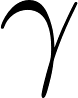<https://www.codecogs.com/eqnedit.php?latex=%5Cgamma#0> | F1 | 0.53 | 0.69 | 0.66 | 0.06 | 0.80 | 4.03 | 2.53 | 0.23 |
| 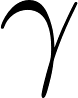<https://www.codecogs.com/eqnedit.php?latex=%5Cgamma#0> | M | 0.65 | 0.77 | 0.51 | 0.08 | 0.52 | 5.00 | 3.54 | 0.56 |
| 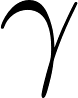<https://www.codecogs.com/eqnedit.php?latex=%5Cgamma#0> | S1 | 0.78 | 0.86 | 0.36 | 0.09 | 0.31 | 6.68 | 5.70 | 1.98 |
| 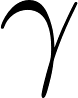<https://www.codecogs.com/eqnedit.php?latex=%5Cgamma#0> | S2 | 0.90 | 0.95 | 0.21 | 0.10 | 0.13 | 10.36 | 13.95 | 12.25 |
| 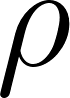<https://www.codecogs.com/eqnedit.php?latex=%5Crho#0> | F2 | 0.40 | 0.60 | 0.90 | 0.06 | 1.13 | 3.32 | 1.96 | -0.19 |
| 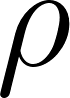<https://www.codecogs.com/eqnedit.php?latex=%5Crho#0> | F1 | 0.53 | 0.69 | 0.75 | 0.08 | 0.75 | 3.88 | 2.56 | -0.29 |
| 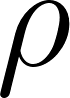<https://www.codecogs.com/eqnedit.php?latex=%5Crho#0> | M | 0.65 | 0.77 | 0.60 | 0.09 | 0.49 | 4.74 | 3.59 | -0.43 |
| 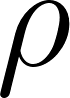<https://www.codecogs.com/eqnedit.php?latex=%5Crho#0> | S1 | 0.78 | 0.86 | 0.45 | 0.10 | 0.28 | 6.17 | 5.82 | -0.65 |
| 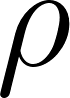<https://www.codecogs.com/eqnedit.php?latex=%5Crho#0> | S2 | 0.90 | 0.95 | 0.30 | 0.11 | 0.11 | 8.98 | 14.59 | -1.13 |
| 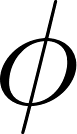<https://www.codecogs.com/eqnedit.php?latex=%5Cphi#0> | F2 | 0.40 | 0.60 | 0.90 | 0.05 | 1.12 | 3.37 | 1.97 | -0.15 |
| 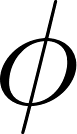<https://www.codecogs.com/eqnedit.php?latex=%5Cphi#0> | F1 | 0.53 | 0.69 | 0.75 | 0.06 | 0.74 | 3.96 | 2.57 | -0.23 |
| 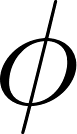<https://www.codecogs.com/eqnedit.php?latex=%5Cphi#0> | M | 0.65 | 0.77 | 0.60 | 0.08 | 0.48 | 4.88 | 3.60 | -0.33 |
| 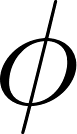<https://www.codecogs.com/eqnedit.php?latex=%5Cphi#0> | S1 | 0.78 | 0.86 | 0.45 | 0.09 | 0.27 | 6.44 | 5.86 | -0.49 |
| 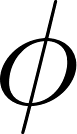<https://www.codecogs.com/eqnedit.php?latex=%5Cphi#0> | S2 | 0.90 | 0.95 | 0.30 | 0.10 | 0.10 | 9.60 | 14.82 | -0.82 |

### **Figure S2**. **Forecasting error is primarily determined by the focal species’ demography and density-dependent pathway, with minimal influence of the competitor’s life history.** Mean absolute percentage error (MAPE) of each predictive model across combinations of focal and competitor life-history strategies and across alternative density-dependent vital rates (juvenile survival, adult survival, progression, retrogression, and fertility). Panels (or groupings) represent distinct species combinations, denoted as “C: F”, where “C” refers to the competitor species and “F” to the focal species; numerical identifiers indicate life-history types positioned along the slow–fast continuum. For each combination, MAPE is averaged across replicate stochastic simulations and across the full range of stage dependence (<https://www.codecogs.com/eqnedit.php?latex=r_%5CDelta#0>) values. The stage-structured model including stage dependence (Model AS) consistently achieved near-zero error, whereas error in the simplified model (Model A) varied depending on the density-dependent vital rate but showed little systematic sensitivity to the life-history strategy of the competitor species.

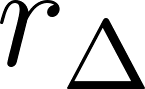

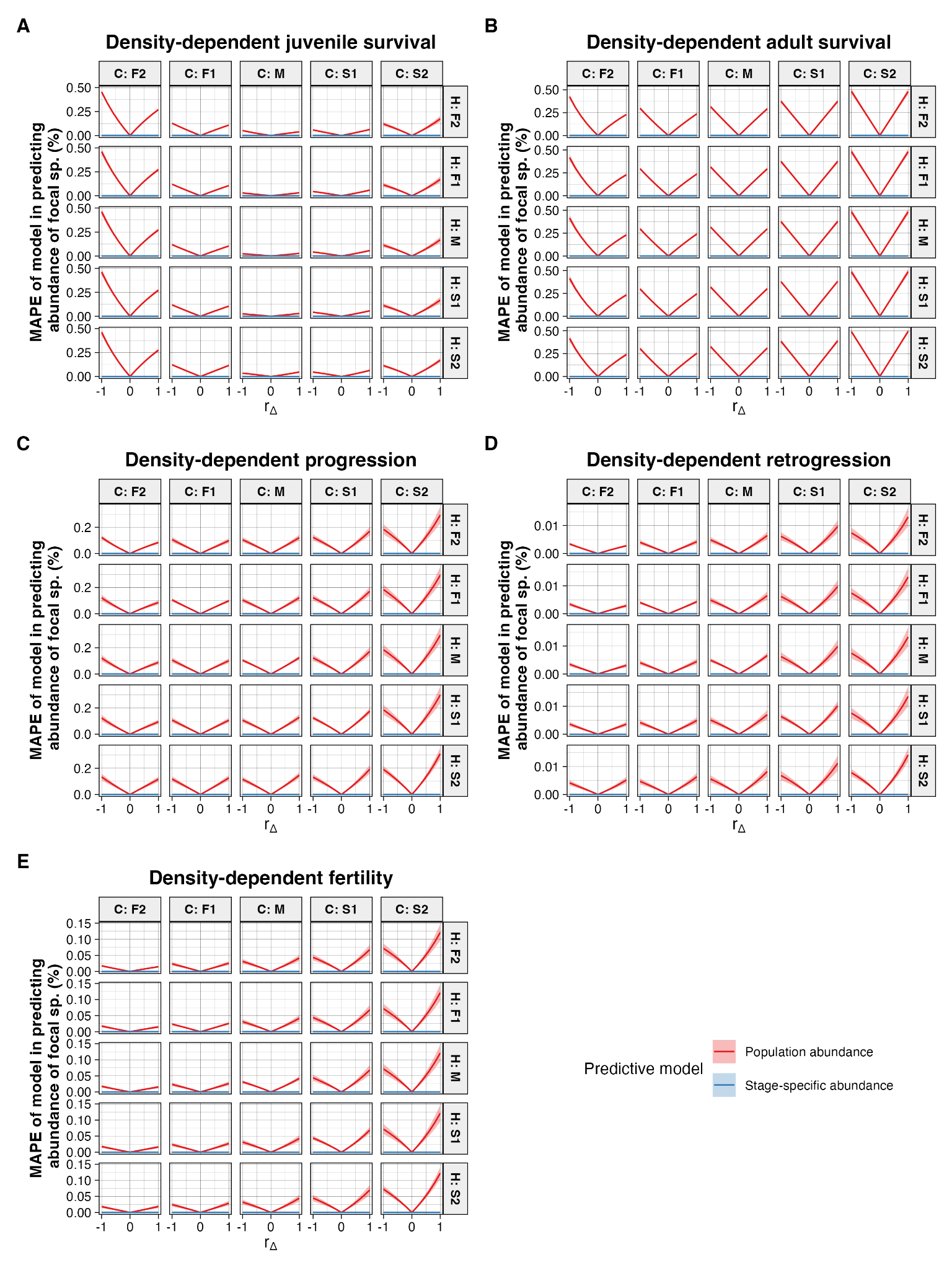

### **Figure S3. Temporal variation in stage structure is low across life histories and strengths of stage dependence.** Standard deviation of the proportion of juveniles in each species across stochastic community simulations, calculated over the full sampling window following initial equilibration. Values are shown for each combination of focal and competitor life-history strategies and across the range of stage dependence (<https://www.codecogs.com/eqnedit.php?latex=r_%5CDelta#0>) values. Species combinations are denoted as “C: F”, where “C” refers to the competitor and “F” to the focal species; numerical identifiers indicate life-history types ordered along the fast-slow continuum. Across all simulations, variability in the juvenile proportion remained small (coefficient of variation < 0.036), with modest differences among life-history strategies. Variation was not systematically amplified by increasing stage asymmetry, indicating that stage composition remained close to its stationary expectation under the imposed stochastic dynamics.

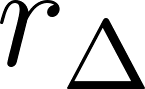

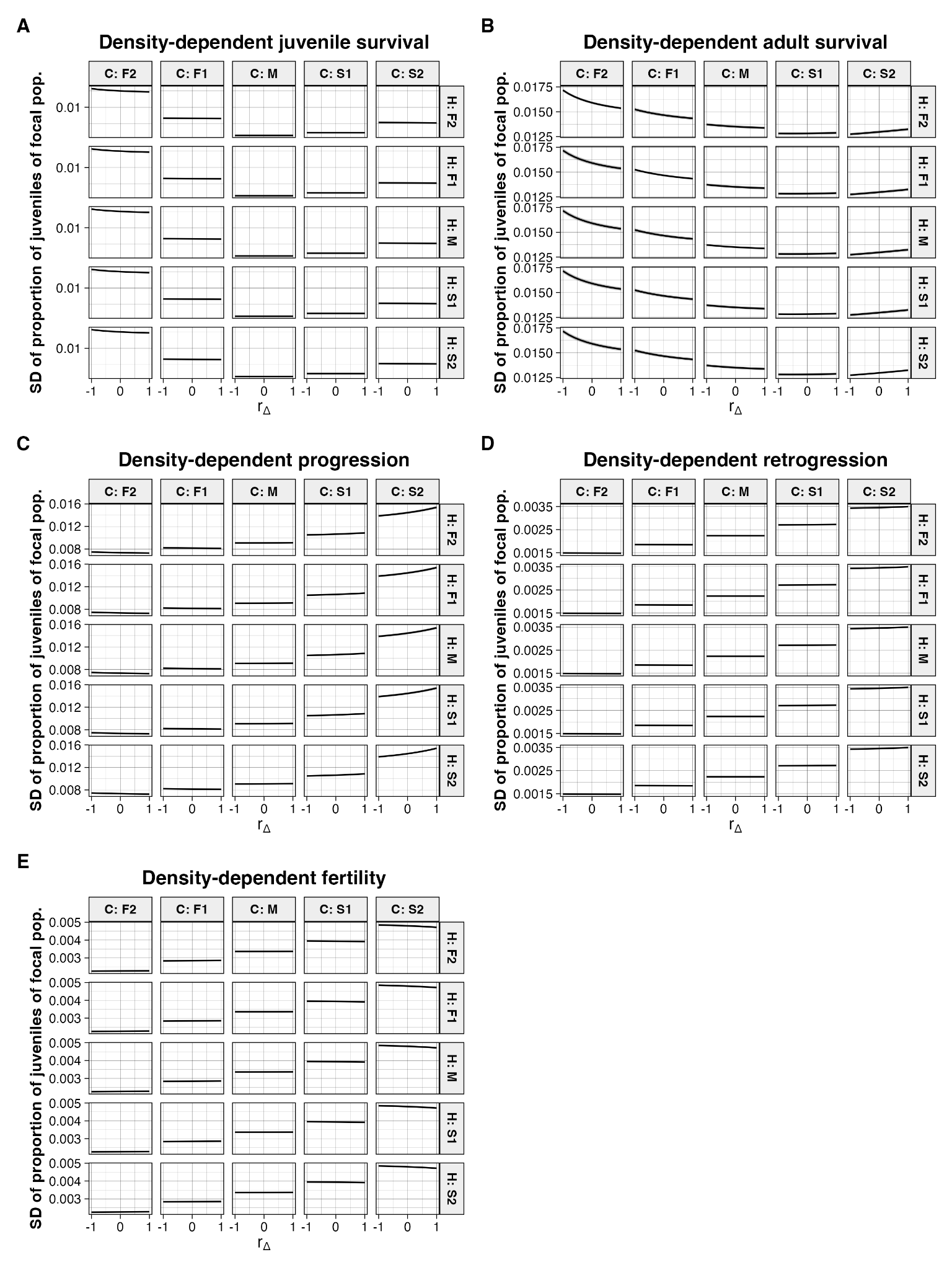
